# Bioactive coatings on 3D printed scaffolds for bone regeneration: Use of Laponite™ to deliver BMP-2 for bone tissue engineering – progression through *in vitro*, chorioallantoic membrane assay and murine subcutaneous model validation

**DOI:** 10.1101/2023.10.25.560313

**Authors:** Karen. M. Marshall, Jonathan P. Wojciechowski, Cécile Echalier, Sebastien J. P. Callens, Tao Yang, Øystein Øvrebø, Vineetha Jayawarna, Janos M. Kanczler, Molly M. Stevens, Jonathan I. Dawson, Richard O.C. Oreffo

## Abstract

Fracture non-union occurs as a consequence of various factors, leading to the development of potentially substantial bone defects. Biomaterial-based approaches for bone regeneration aim to explore alternative strategies to repair non-healing fractures and critical-sized bone defects. Thus, rigorous assessment of the ability to translate biomaterials towards clinical use is vital. Growth factors induce an effect on cells to change their phenotype, behaviour and initiate signalling pathways, leading to an effect on matrix deposition and tissue formation. Bone morphogenetic protein-2 (BMP-2) is a potent osteogenic growth factor, with a rapid clearance time *in vivo* necessitating clinical use of high doses, with potential deleterious side-effects. This work explored the potential for Laponite™ nanoclay coating of poly(caprolactone) trimethacrylate (PCL-TMA900) scaffolds to bind BMP-2 for enhanced osteoinduction. *In vitro* experiments confirmed the cytocompatibility of the PCL-TMA900 scaffolds and effective osteogenic differentiation of C2C12 myoblast cells in response to the Laponite/BMP-2 coating. The chorioallantoic membrane (CAM) assay verified PCL-TMA900 scaffold material biocompatibility and ability to support angiogenesis. A murine subcutaneous implantation model assessed heterotopic bone formation in response to the Laponite/BMP-2 coating, when used immediately post-coating and after 24 hours of room temperature storage, to evaluate a delayed use manner. The Laponite/BMP-2 coated PCL-TMA900 scaffolds implanted showed consistent, significant bone formation over the study period compared to the uncoated PCL-TMA 900 scaffold and BMP-2 only coated control scaffolds *in vivo*, indicating the ability of Laponite to bind the BMP-2 to the PCL-TMA900 scaffold. Bone formed peripherally around the Laponite/BMP-2 coated scaffold, with no aberrant bone formation observed. The Laponite/BMP-2 coating was found to retain its bioactivity after storage for 24 hours prior to use *in vivo*, however this was not to the same volume or reliability of bone formation as when used immediately post-coating. To take these studies forward, the Laponite/BMP-2 coating warrants examination in a critical-sized bone defect model to assess efficacy in an osseous site.

## 1. Introduction

Due to advances in healthcare, global longevity is increasing, creating new challenges including the prevalence of chronic disease and age-related tissue degeneration which can fail to heal after injury, such as bone following a fracture. Bone defects can arise from a variety of causes and in severe cases, amputation may be required due to extensive soft tissue, vascular and/or nerve damage, making the limb unsalvageable [1, 2]. Whilst tissue grafts are the current gold standard, grafting tissue has inherent risks, including the potential for immune-mediated rejection or infection. Bone tissue engineering seeks to develop materials to substitute the use of patient bone, (i.e., autografts, allografts) to encourage rapid healing of large bone defects [3, 4]. Advances in tissue engineering have enabled the development of materials that mimic the composition and functionality of the damaged tissue, thereby reducing the need for grafting or transplantation procedures. However, bone tissue is a complex tissue to recapitulate, hence, the ideal bone substitute material is biocompatible, biodegradable, osteoconductive, osteoinductive, porous, strong, easy to use and cost effective [1].

In general, bone morphogenetic protein-2 (BMP-2) is commonly administered at a high concentration on materials to elicit a biological response (i.e., µg/mL in small animal studies and mg/mL in large animal studies and clinically in humans). These high concentrations of BMP-2 may saturate the binding sites of the scaffold and result in undesirable burst release of this potent growth factor when implanted [5]. Therefore, to maintain bioactivity whilst reducing and minimising potential side-effects, BMP-2 has been immobilised onto scaffolds by various methods including the use of an avidin-biotin binding system, coating with fibronectin or heparin and within nanoparticles, hydrogels, or microspheres [6–12]. In other examples, BMP-2 and vascular endothelial growth factor (VEGF) bound to a scaffold by antibodies against each growth factor respectively, led to osteogenic and angiogenic responses of bone marrow stromal cells and biocompatibility in the chorioallantoic membrane (CAM) assay [13]. Additionally, coating coralline hydroxyapatite granules with octa-calcium phosphate and BMP-2 produced a biocompatible construct. It was observed that BMP-2 incorporated into the coralline hydroxyapatite coating provided superior results compared to BMP-2 simply adsorbed onto the scaffold surface [14]. Covalent approaches such as the application of a chemical deposition technique to coat titanium with glycidyl methacrylate (GMA) to bind rhBMP-2 provided high results of rhBMP-2 bound to the titanium substrates (243.9 ± 25.7 ng), compared to physical adsorption (40.2 ± 32.7 ng), however the BMP-2 adsorbed to the surface of the titanium was not compared to the mass of BMP-2 bound via GMA with regard to osteogenic differentiation of cells upon the scaffold [15]. To mitigate and avoid adverse effects and maintain the BMP-2 *in situ* to induce bone formation, the use of a low concentration/mass of BMP-2 adhered to poly(ethyl acrylate) and fibronectin coated materials has been applied by the Salmeron-Sanchez group, with bone healing demonstrated in a murine radial defect study as well as in veterinary clinical cases [7, 16]. Despite these methods of immobilised delivery of growth factors to tissues, further optimisation is required to ensure the growth factor is active and at the correct concentration and mass to produce a biological effect.

The current work describes a 3D-printed octet-truss scaffold design using an optimised formulation of PCL trimethacrylate (PCL-TMA900). This polymer permitted the printing of intricate shapes with less risk of fragmentation, as the polymer was less hard and brittle than the PCL trimethacrylate (PCL-TMA) used in our previous study [17]. The objective of the current study was to investigate a biocompatible and biodegradable smectite nanoclay, Laponite™, previously demonstrated to sequester growth factors, as a dry coating on the PCL-TMA900 scaffold, to adhere bone morphogenetic protein-2 (BMP-2) to the PCL-TMA 900 scaffold and improve its osteogenic potential [18–20].

Laponite (Na_0.7_Si_g_Mg_5.5_Li_0.3_O_20_(OH)_4_) has injectable properties due to the shear stress on the particles, which coalesce to form a 3D structure making Laponite thixotropic. The gel can be modified to become denser by application of protein or salt-containing solutions. Due to these beneficial properties, Laponite has been studied extensively as a tissue regenerative biomaterial [21]. Laponite gels can be made in a range of concentrations, with 3% (w/w) and above leading to a ‘house of cards’ structure of the charged particle discs, producing a stiffer gel [21]. Laponite has previously been used as a coating on decellularized bone to sequester BMP-2 from culture media, leading to C2C12 myoblast cell osteogenic differentiation on the bone surface [18].

The study set out to investigate the use of Laponite/BMP-2 coatings as a dry, ready-to-use, biocompatible, bioactive, reliable, osteoinductive coating on PCL-TMA900 scaffolds. This relied upon retaining active BMP-2 on or within the Laponite surrounding the PCL-TMA900 scaffold in sufficient quantities to induce osteogenic differentiation of skeletal cell populations and to produce significant bone volumes *in vivo.* Given the biocompatibility of Laponite and BMP-2 are known, the Laponite/BMP-2 coating could easily and practically be applied to induce bone formation around the PCL-TMA900 scaffold *in vivo*. The PCL-TMA900 was examined for cytocompatibility with human bone marrow stromal cells (HBMSCs) using alamarBlue™ cell viability reagent and biocompatibility using the CAM assay, compared to the PCL-TMA scaffold material previously studied [17]. Ethidium homodimer staining was used to visualise the low viscosity Laponite (1% w/w) covering the PCL-TMA900 scaffold surface. The ability for Laponite to remain adherent to the PCL-TMA900 scaffold material was assessed and fluorescent fluorescein isothiocyanate labelled bovine serum albumin (FITC-BSA) was employed to visualise the labelled protein on the Laponite coated PCL-TMA900 scaffolds. Subsequently, it was necessary to ensure BMP-2 had bound to the Laponite and dry BMP-2 could retain its bioactivity, evidenced by ALP staining on C2C12 cell seeded scaffolds. Furthermore, the optimal incubation time and concentration of BMP-2 solution applied to the Laponite coating was investigated. After *in vitro* confirmation of bioactivity, *in vivo* murine subcutaneous implantation studies were completed to assess the efficacy of the Laponite/BMP-2 coating on the PCL-TMA900 scaffolds to induce bone formation, prior to future use in bone defect orthotopic models.

## 2. Materials and Methods

### 2.1 Materials

Reagents were purchased as follows: collagenase (Gibco, UK); recombinant human BMP-2 (Infuse/InductOS® Bone graft kit, Medtronic, USA); BMP-2 Quantikine ELISA Kit (biotechne®, R&D systems, UK); alcian blue 8X, light green SF, orange G 85% pure, paraformaldehyde 96% extra pure, phosphomolybdic acid hydrate 80% (Acros Organics); picrosirius Red, Van Gieson’s stain, Weigert’s Haematoxylin Parts 1 and 2 (Clintech Ltd, UK); Phosphate Buffered Saline (PBS), trypsin/ethylenediaminetetraacetic acid (EDTA), Dulbecco’s Modified Eagle Medium (DMEM), Alpha Minimum Essential Medium (αMEM), penicillin-streptomycin (P/S) (Scientific Laboratory Supplies, SLS); acetic acid, acetone, acid fushsin, alizarin red S, fast violet B salts, histowax, hydrochloric acid, Naphthol AS-MX phosphate 0.25%, parafilm, ponceau xylidine, silver nitrate, sodium hydroxide pellets (Merck, UK); alamarBlue™ HS Cell Viability Reagent, 70 µm cell strainer, dibutyl phthalate xylene (DPX), ethidium homodimer-1, fetal calf serum (FCS), fisherbrand grade 01 cellulose general purpose filter paper, Histoclear, Vybrant™ CFDA SE Cell Tracer Kit (Thermofisher Scientific, UK); fast green and sodium thiosulphate (VWR); Lubrithal (Dechra, UK), Isoflurane (Dechra, UK), Buprenorphine (Buprecare® multidose, Animalcare, UK) and Vetasept® sourced from MWI animal health, UK. 5/0 PDS II suture (Ethicon, USA) from NHS supply chain. All other consumables and reagents were purchased from Sigma-Aldrich, UK.

### 2.2 Production of PCL trimethacrylate scaffold material

#### 2.2.1 Poly(caprolactone) trimethacrylate synthesis – low and high molecular weight (PCL-TMA and PCL-TMA900)

PCL-TMA (poly(caprolactone) triol, trimethylolpropane initiated, M_w_ = 300 Da) has been synthesised and 3D printed via stereolithography (SLA) previously [17, 22–24] (**Supplementary information Figure 1**). Poly(caprolactone) triol, trimethylolpropane initiated, M_w_ = 830 Da was purchased from BOC Sciences.

To synthesise the PCL-TMA900 material, poly(caprolactone) triol, M_n_ = 830 Da, (100 g, 0.12 mmol, 1 eq), anhydrous dichloromethane (300 mL) and triethylamine (100 mL, 0.72 mmol, 6 eq) were added to a 1 L two-necked round bottom flask. The reaction was placed under a nitrogen atmosphere and then cooled in an ice-water bath for 15 minutes. A pressure-equalising dropper funnel charged with methacryloyl chloride (53 mL, 0.67 mmol, 4 eq) was attached to the round bottom flask. The methacryloyl chloride was added dropwise over approximately 3 hours. The reaction was covered with aluminium foil to protect it from light and allowed to stir and warm to room temperature (RT) overnight. The next day, methanol (50 mL) was added to quench the reaction, which was allowed to stir at RT for 30 minutes. The reaction mixture was dissolved in dichloromethane (800 mL), transferred to a separating funnel and washed with 1 M aqueous hydrochloric acid solution (5×250 mL), saturated sodium bicarbonate solution (4×400 mL) and brine (1×400 mL). The organic layer was then dried with anhydrous magnesium sulphate, filtered and concentrated via rotary evaporation. The crude yellow liquid was then purified using a silica plug, with dichloromethane as the eluent. Fractions containing PCL-trimethacrylate were pooled and concentrated via rotary evaporation. The PCL-trimethacrylate was transferred to a brown glass vial and dried using a stream of air (through a plug of CaCl_2_) overnight to yield the title compound as a slightly yellow viscous liquid (94.25 g). The PCL-trimethacrylate was supplemented with 200 ppm (w/w) of 4-methoxyphenol (MEHQ) as an inhibitor.

^1^H NMR (400 MHz, CDCl_3_) δ 6.11 – 6.05 (m, 3H), 5.61 – 5.50 (m, 3H), 4.16 – 3.99 (m, 18H), 2.34-2.27 (m, 12H), 1.96 – 1.88 (m, 9H), 1.71 – 1.60 (m, 26H), 1.47 – 1.31 (m, 12H), 0.98 – 0.83 (m, 3H).

The characterisation data agreed with that previously reported [22]. The degree of functionalisation is determined as >90% (**Supplementary information Figure 2**).

#### 2.2.2 3D printing of PCL-TMA and PCL-TMA900

The PCL-TMA and PCL-TMA900 octetruss scaffold shape was modelled in Rhinoceros 3D software (McNeel, Europe, Barcelona, Spain) using a diameter and height of 5 mm, strut thickness of 1 mm, surface area 143.4 mm^2^.

The PCL-TMA and PCL-TMA900 scaffolds were printed using masked SLA 3D printing on a Prusa SL1. The resin was prepared for 3D printing by first dissolving 0.1% (w/w) 2,5-thiophenediylbis (5-tert-butyl-1,3-benzoxazole) (OB+) as a photoabsorber in the PCL trimethacrylate by stirring at RT for 1 hour. Finally, 1.0% (w/w) diphenyl(2,4,6-trimethylbenzoyl)phosphine oxide (TPO-L) as a photoinitiator was added to the resin.

After printing, the scaffolds were rinsed with 100% ethanol and removed from the build plate. Scaffolds were sonicated in ethanol (5×5 minutes) and allowed to dry for 15 minutes at RT. The scaffolds were post-cured using a Formlabs Form Cure for 60 minutes at RT. After post-curing, the scaffolds were soaked into ethanol overnight at RT on a rocker (100 rpm), rinsed with 100% ethanol (3×) and allowed to dry at RT before being EO sterilised.

#### 2.2.3 Compressive testing of PCL-TMA and PCL-TMA900

The compressive mechanical properties of the scaffolds were measured using a Bose Electroforce TA 3200 equipped with uniaxial load cells. The compressive properties of PCL-TMA900 were assessed using ISO 604 standards, with 10 × 10 × 4 mm (H × W × D) samples printed as described above. The samples were tested with the layers printed perpendicular and parallel or the strain applied. Due to the geometry of the unit cell scaffolds (PCL-TMA and PCL-TMA900), the effective elastic modulus was measured. A 1 N preload was implemented, with the elastic modulus of the PCL-TMA900 ISO 604 standards measured between 2 – 4% strain (n = 5) and the effective elastic modulus of the PCL-TMA and PCL-TMA900 unit cells measured between 1 – 3% strain (n = 5) (**Supplementary information Figures 3 and 4**)

### 2.3 HBMSC isolation and culture for cytocompatibility assay

#### 2.3.1 Isolation and culture of HBMSCs

The method is extrapolated from Marshall *et al*. [17]. Briefly, human bone marrow was collected from patients, identified by sex (male (M) or Female (F)) and age (e.g., F60) only, to maintain confidentiality, with approval of the University of Southampton’s Ethics and Research Governance Office and the North-West Greater Manchester East Research Ethics Committee (18/NW/0231). In a class II hood, 5- 10 mL alpha-Minimum Essential Medium (α-MEM, Lonza, UK) or Dulbecco’s Modified Eagle Medium (DMEM, Lonza, UK) was used to wash the marrow repeatedly until any bone was white, no fat/blood material remained, and the sample tube was shaken vigorously to extract the HBMSCs. The supernatant media/cellular debris mix was collected in a 50 mL falcon tube. The cell suspension was centrifuged (272g Heraeus mega 1.0R centrifuge) for 5 minutes. The supernatant was removed, the cell pellet resuspended in α-MEM or DMEM and passed through a 70 µm cell strainer to remove bone and fat debris. The cell suspension was centrifuged again, and the supernatant poured off. The cell pellet was resuspended in basal media (α-MEM, 10% fetal calf serum (FCS), 1% penicillin-streptomycin (P/S)) and the HBMSCs were cultured in T175 flasks at 37 °C in 5% CO_2_/balanced air until approximately 80% confluent. Media was changed every 3-4 days. Collagenase (2% solution and or 0.22 IU/mg) was applied to the cell culture prior to the trypsin solution (1x concentration (Stock Trypsin/EDTA (10X), includes 1,700,000 U\L trypsin 1:250 and 2g/L Versene® (EDTA)), for passaging and seeding onto PCL-TMA and PCL-TMA900 scaffolds.

#### 2.3.2 Seeding HBMSCs onto scaffolds

2.5×10^4^ Passage 2 (P2) HBMSCs were seeded per scaffold, set up in triplicate, for cell viability/alamarBlue™ HS experiments as previously described [17]. PCL-TMA and PCL-TMA900 octetruss scaffolds were sterilised with 70% ethanol, rinsed twice in phosphate buffered saline (PBS) and then twice in α-MEM and added individually to 2 mL Eppendorf tubes with 500 µL cell suspension (α-MEM, 1% P/S, without FCS). The Eppendorf tubes were placed in a 50 mL falcon tube positioned horizontally on the MACSmix™ Tube Rotator (6 eppendorf tubes per 50 mL tube and 4 x 50mL tubes per rotator). The Eppendorf tubes were fully sealed and therefore the only air available to cells was within each tube, however, the media maintained its normal cell culture colour, indicating the pH of the media was unaltered. FCS was added at approximately 10% (55 µL) to each tube after 3 hours. After 24 hours, each scaffold was individually cultured in 1.5 mL basal media in a 24 well plate. Culture was performed at 37 °C in 5% CO_2_/balanced air, with media changes every 3 days.

### 2.4 alamarBlue™ HS Cell Viability assay

The method has been previously described [17]. In brief, for cell viability experiments, alamarBlue™ HS Cell Viability Reagent was added to basal media at 10% (v/v) concentration. Fluorescence measurements were taken at day 1 (when the PCL-TMA/PCL-TMA900 scaffolds were removed from the Eppendorf tube after 24 hours of cell seeding) and on day 14 (each scaffold was moved to a new 24 well plate to ensure only adherent cells were quantified). A 1 mL aliquot of media/alamarBlue™ HS mix was added to each well and to three wells with no scaffold as background measurements and incubated for 4 hours at 37 °C in 5% CO_2_/balanced air. After 4 hours, 100 µL samples were taken from each well and plated in triplicate in a black 96 well plate. Fluorescence was measured using the GloMax® Discover Microplate Reader (Promega, UK) at Green 520 nm excitation and 580-640 nm emission and the average background measurement was subtracted from each sample well value. The increase in the number of cells was proportional to the increase in measured fluorescence, correlating cell number to fluorescence (ref 1^st^ paper supplementary information).

### 2.5 Live/dead labelling of HBMSCs on PCL-TMA and PCL-TMA900 scaffolds

The method has been previously described [17]. In brief, Vybrant Cell Tracer 10 μM and 5 µg/mL Ethidium homodimer-1 in PBS were used to label live and dead cells respectively, on PCL-TMA and PCL-TMA900 scaffolds at day 1 and day 14 post alamarBlue™ HS analysis. Media/alamarBlue™ HS was removed and the scaffolds washed twice in PBS. 1 mL labelling solution was added to cover the scaffolds and incubated for 15 minutes at 37 °C in 5% CO_2_/balanced air. The labelling solution was removed and replaced with α-MEM/1% P/S/10% FCS and culture continued for 30 minutes. The samples were imaged with fluorescence microscopy using the FITC filter (excitation 485/20 nm, emission 515 nm) for live cells and RHODA/TRITC filter (excitation 510-560 nm, emission 590 nm) for dead cells, with a Zeiss Axiovert 200 microscope and Axiovision 4.2 imaging software.

### 2.6 Laponite (1% w/w) production

1% (w/w) Laponite was made 48 hours prior to use by adding 0.1 g Laponite (BYK-Altana) to 9.9 g dH_2_O very slowly in a glass jar, while mixing at 250 rpm with a magnetic stirrer at RT and left to stir for 1-2 hours to ensure complete dissolution. The jar was weighed and autoclaved at 126 °C in a bench-top autoclave and reweighed. In a sterile hood, the mass of dH_2_O lost was replaced by adding sterilised dH_2_O until the original mass was reached and stored at 4 °C.

### 2.7 Ethidium homodimer staining for Laponite presence and persistence on the PCL-TMA900 scaffolds

In a sterile hood, PCL-TMA900 scaffolds were immersed in Laponite or dH_2_O for 1 hour at RT, removed from the liquid and allowed to dry, suspended by sterile 2/0 nylon suture material (Ethicon) on sterile autoclaved paper/plastic coated cups without touching any surface (**Supplementary information Figure 5**) and stored in a 50 mL falcon tube at 4 °C. The dH_2_O scaffolds are denoted as uncoated controls as the dH_2_O evaporated. The scaffolds were placed in PBS 7 days, 2 days, 1 day and 1 hour prior to ethidium homodimer staining, to visualise red colouration of Laponite on the scaffolds. A 1 mL aliquot of 5 µg/mL ethidium homodimer in PBS was added to each scaffold condition (n = 3) for 15 minutes prior to imaging with a digital camera (Canon G10) attached to a stemi 2000-c stereomicroscope (Zeiss, UK). The images were enhanced by increasing the colour saturation to 200% for clearer illustration of the red colour on the scaffold surface.

### 2.8 FITC-BSA coating of scaffolds to assess protein binding by Laponite

PCL-TMA900 scaffolds (uncoated or 1% Laponite coated) were immersed for 1 hour at RT in PBS, bovine serum albumin (BSA) diluted in PBS (50 µg/mL or 100 µg/mL in PBS), or Fluorescein isothiocyanate-BSA (FITC-BSA) diluted in PBS (50 µg/mL or 100 µg/mL) and imaged by fluorescence microscopy using the FITC filter (excitation 485 nm, emission 515 nm) on the Zeiss Axiovert 200 microscope and Axiovision 4.2 imaging software (Zeiss, UK).

### 2.9 Laponite and BMP-2 coating of PCL-TMA900 scaffolds and C2C12 cell seeding to assess differentiation by visualisation of ALP staining

Two PCL-TMA900 octetruss-type scaffolds per 2 mL eppendorf containing 1 mL of 1% Laponite or 1 mL of sterile dH_2_O were immersed for 1 hour at RT. The scaffolds were removed, suspended and left to dry at RT for 2-3 hours (**Supplementary information Figure 5**). The dH_2_O scaffolds are denoted as uncoated as the dH_2_O evaporated. The dry Laponite coated or uncoated scaffolds were individually submerged in 1 mL of BMP-2 solution of 5 µg/mL or 1 µg/mL (1.5 mg/mL BMP-2 diluted in PBS) in 1.5 mL low-protein binding eppendorf tubes for 24 hours at RT and BMP-2 solution (5 µg/mL and 1 µg/mL) without a scaffold was used as a control for the ELISA. The resultant scaffolds were therefore Laponite/BMP-2 coated or BMP-2 only coated. Each scaffold was removed and placed individually in a well of a 24 well plate to dry for 2-3 hours, rather than suspending it. Once dry, the scaffolds were individually placed in 2 mL eppendorf tubes for C2C12 cell seeding on the rotator. The remaining BMP-2 solutions were stored at 4 °C for use in an ELISA. The optimal incubation time and concentration of BMP-2 solution applied to the Laponite coating was investigated (**Supplementary information Figure 6**).

Each scaffold was seeded with 2×10^5^ C2C12 (P21) cells in 500 µL of DMEM/1% P/S/10% FCS (basal) media within a 2 mL eppendorf using the MACSmix™ magnetic rotator propped up on a tip box to keep the scaffolds submerged for 24 hours. BMP-2 at 5 µg/mL was added to the positive controls. The negative controls had no BMP-2 on the scaffolds or in the media. The remaining scaffolds were cultured in basal media to allow the absorbed/attached BMP-2 only to be assessed. After 24 hours, the scaffolds seeded with C2C12 cells were moved to a 24 well plate and 1.5 mL of media was added (basal with 2% FCS ± 5 µg of BMP-2 (3.33 µg/mL BMP-2). Culture was continued for 4 days and ALP staining performed on the 5^th^ day from cell seeding. Ethidium homodimer staining (5 µL/mL PBS) was used to label the cell nuclei to visualise the C2C12 cell confluency on the Laponite/BMP-2 coated and BMP-2 only coated PCL-TMA900 scaffolds. Images were taken using the RHODA/TRITC filter (excitation 510-560 nm, emission 590 nm) with the Zeiss Axiovert 200 microscope and Axiovision 4.2 imaging software (Zeiss, UK).

### 2.10 ALP staining of cells on tissue culture plastic (TCP) and scaffolds

Media was removed and the scaffolds/cells were washed twice in PBS followed by addition of 90% ethanol for 10 minutes, prior to washing again in PBS. Alkaline phosphatase activity was visualised by creating a staining solution of 0.1 mg/mL Naphthol AS-MX phosphate (Sigma) and 0.0024% (w/v) Fast Violet-B salt (Sigma) mixed in 3.0285g Tris/800 µL 100% hydrochloric acid/250 mL dH_2_O (pH 9) and adding 1000 µL or 300 µL staining solution to scaffolds or cells on tissue culture plastic respectively. Scaffolds/cells were incubated in the dark at 37 °C for 1 hour. Red stain indicated positive ALP activity. dH_2_O was added to stop the reaction and the solution removed prior to imaging using a digital camera (Canon G10) attached to a stemi 2000-c stereomicroscope (Zeiss, UK).

### 2.11 BMP-2 ELISA for quantification of BMP-2 remaining after PCL-TMA900 scaffold coating for in vitro and in vivo experiments

The Quantikine® ELISA BMP-2 immunoassay kit was used to test the BMP-2 remaining in the scaffold coating solution, to deduce the quantity of BMP-2 bound to the scaffold. Reagents and standard solutions were prepared as per kit instructions. The BMP-2 solutions were diluted in calibration buffer to enable results to be extrapolated from the standard curve. A 100 µL aliquot of assay diluent followed by 50 µL of standard, control, or sample was added to each well. The plate was shaken on a horizontal orbital microplate shaker at 450 rpm for 2 hours at RT. The solution was aspirated, and the wells washed and aspirated again four times. BMP-2 conjugate (200 µL) was added to each well and the incubation step on the shaker repeated for 2 hours. The aspiration and wash steps were repeated and 200 µL of substrate solution added to each well and the plate was left for 30 minutes at RT in the dark on the benchtop. Finally, 50 µL of stop solution was added to each well and the colour changed from blue to yellow. The plate was read using a GloMax® Discover microplate reader set to 450 nm. Results were calculated from the standard curve to determine the concentration of BMP-2 in each sample. The 5 µg/mL and 1 µg/mL stock coating solutions (with no scaffold immersed) were n = 1, which were plated out in triplicate, while the 5 µg/mL or 1 µg/mL BMP-2 solutions used to coat BMP-2 only and Laponite coated scaffold samples were n = 3 and plated out once each as biological triplicates. The *in vivo* subcutaneous implantation study samples were n = 4 biological samples for each scaffold type (Laponite/BMP-2 or BMP-2 only coated) with each sample plated out in triplicate wells and the average result of each scaffold triplicate was used for statistical analysis.

### 2.12 Laponite and BMP-2 coating of a 24 well plate to assess the osteogenic response of C2C12 cells to delayed use of the coating

A 300 µL aliquot of 1% Laponite was applied in three wells of a 24 well plate and allowed to dry. Then, a 1 mL aliquot of 5 µg/mL BMP-2 solution was added for 24 hours. After 24 hours, the remaining solution was removed and the wells allowed to dry and this process was repeated for another three wells. In addition, Laponite was added to six wells and allowed to dry for the control wells. Of the six Laponite coated wells, three had no BMP-2 added (negative controls) and three had BMP-2 (200 ng/mL) added to the media (positive controls) at cell seeding. 5×10^4^ cells C2C12 cells (P15) were seeded per well in basal media (DMEM/1%P/S/10%FCS) or basal media containing BMP-2 (200 ng/mL). The media was not changed and ALP staining was performed after 3 days.

### 2.13 Statistical analysis

alamarBlue™ results were set up with biological triplicates, with triplicate readings taken from each sample and the average of the triplicate readings used for statistical analysis. Analysis and graphical presentation were performed using GraphPad Prism 9, version 9.2.0. P values <0.05 were considered significant. Graphical representation of significance as follows: ns is no significant difference, *p<0.05, **p<0.01, ***p<0.001, ****p<0.0001. All data presented as mean and standard deviation (S.D.).

### 2.14 Chorioallantoic membrane assay

Descriptions are extrapolated from Marshall *et al.* [25]. All procedures were performed with prior received ethical approval and in accordance with the guidelines and regulations stated in the Animals (Scientific Procedures) Act 1986, however a UK Personal Project License (PPL) was not required as the eggs were only used until embryonic day (ED) 14. Briefly, fertilised hens (Gallus gallus domesticus) eggs (Medeggs, Henry Stewart & Co., Lincolnshire) were incubated in a humidified (60%), warm (37 °C) incubator (Hatchmaster incubator, Brinsea, UK) at ED 0. The eggs were incubated in a horizontal position, on a rotating pattern (1 hour scheduled rotation), prior to albumin removal at ED 3 ([17] supplementary information). The minimum number of eggs required to see a significant difference (p<0.05) between groups was calculated with power 80%, with n=7 used in this study in case of any undeveloped eggs.

#### 2.14.1 Scaffold preparation and implantation of materials

PCL-TMA and PCL-TMA900 uncoated scaffolds were sterilised with 70% ethanol followed by rinsing four times with PBS and implanted at ED 7. Descriptions are extrapolated from Marshall *et al.* [25] and [17]. In brief, eggs were candled to check viability and a No. 10 scalpel blade was used to create a 0.5 cm by 0.5 cm window. The white inner shell membrane was peeled away and one scaffold placed onto each CAM. Parafilm covered the window and was secured by labelled tape. The eggs were placed horizontally within the egg incubator at 37 °C and 60% humidity without rotation.

#### 2.14.2 Analysis of results

Samples were harvested at ED 14 of incubation with methodology extrapolated from Marshall *et al*.[25]. Blinding of the assessor was performed. The window was opened digitally and with forceps to image the scaffold/CAM using a stemi 2000-c stereomicroscope (Zeiss, UK) and digital camera (Canon G10) for illustrative, recording purposes only. Quantification of angiogenesis was performed using the Chalkley eyepiece graticule. Three separate counts were made and the average score was calculated for each egg. Biocompatibility was assessed by counting viable, developed chicks and any dead or deformed chicks. Thereafter, the scaffold and surrounding CAM tissue was removed for histological assessment. The scaffold material and adherent tissue were placed into 2 mL 4% PFA in a 24 well plate for 24 hours at 4 °C followed by 70% ethanol for further histological processing. The chick was euthanised by an approved schedule 1 method at ED 14. Data was analysed using GraphPad Prism 9, software version 9.2.0. Data presented as the mean and standard deviation (S.D.). P values <0.05 were deemed significant. Due to non-forming eggs the mean and standard deviation of the Chalkley score results from formed/viable eggs or those able to be counted were included.

### 2.15 *In vivo* subcutaneous implantation study

#### 2.15.1 Mouse type and housing

All procedures were performed in accordance with institutional guidelines, with ethical approval and under PPL P96B16FBD and PPL PP8873619, in accordance with the regulations in the Animals (Scientific Procedures) Act 1986 and using the ARRIVE guidelines. Four, adult, male MF-1 wild type mice, bred on site and group housed in individually ventilated cages (IVCs) were used in each study. Mice had access to *ad libitum* standard pellets and water.

#### 2.15.2 Scaffold preparation

Uncoated (no immersion in any dH_2_O or solutions) PCL-TMA900 scaffolds were the negative control. Two scaffolds per 2 mL eppendorf were immersed in 1 mL dH_2_O or 1 mL 1% Laponite and suspended by suture material in the sterile hood for 2-3 hours followed by immersion individually in BMP-2/PBS solution (5 µg/mL) in a 1.5 mL low protein binding eppendorf for 24 hours, therefore a maximum of 5 µg of BMP-2 could adhere to each scaffold. The dH_2_O scaffolds therefore became uncoated as the dH_2_O evaporated in the first step, prior to immersion in BMP-2 solution, therefore the resultant scaffolds were denoted BMP-2 only coated. The Laponite/BMP-2 coated or BMP-2 only coated scaffolds were removed and left to dry for 2-3 hours in the well of a 24 well plate prior to immediate use or Laponite/BMP-2 coated PCL-TMA900 scaffolds were stored in the plate at RT for ‘delayed’ use after 24 hours. Sterile collagen sponge (4 mm diameter, approximately 2 mm height) soaked with 5 µg of BMP-2 within 15 µL of InductOs® buffer solution (333.33 µg/mL) immediately prior to use provided the positive control (**Supplementary Information Table 3** for buffer constituents). Each scaffold type was n = 4 per group (Uncoated, BMP-2 only (or Delayed Laponite/BMP-2 in the second study), Laponite/BMP-2 coated or collagen sponge/BMP-2). However, in the ‘delayed use’ study, mouse 4 had a ‘delayed’ scaffold implanted which was coated with Laponite/BMP-2 6 months rather than 24 hours previously. Also, the BMP-2 solution applied to the collagen sponge in the ‘delayed use’ study was residual from the first subcutaneous implantation study, kept at 4 °C in a low protein binding eppendorf tube. An ELISA was performed using a Quantikine® ELISA BMP-2 immunoassay kit on the BMP-2 solutions remaining (1:2500 (v/v) dilution by serial dilution) from the coating process of the initial Laponite/BMP-2 and BMP-2 only coated PCL-TMA900 scaffold subcutaneous implantation study to determine the quantity of BMP-2 adhered to each PCL-TMA900 scaffold.

#### 2.15.3 Subcutaneous implantation surgical procedure

The method is extrapolated from Marshall *et al.* [17]. In brief, general anaesthesia was induced using an induction chamber with volatile Isoflurane (5%) and 100% oxygen (0.8 L/min flow rate) and continued for a surgical plane of anaesthesia maintained at approximately 1.5-2.5% isoflurane. Once anaesthetised, Lubrithal was applied to the eyes, ear marking performed for identification, the dorsum of the mouse was shaved and chlorhexidine/alcohol solution (Vetasept® Clear Spray) applied to the dorsum and allowed to dry. Buprenorphine (0.05 mg/kg) was administered subcutaneously. Four dorsal 5 mm incisions were made (2 on each side over the shoulder and flank regions) and the skin elevated using blunt dissection, creating a pouch for each scaffold or collagen sponge/BMP-2 implanted. The skin incisions were closed in a simple interrupted, horizontal mattress suture pattern with absorbable 5/0 PDS (Ethicon), with the knots towards midline. The mice were recovered inhaling 100% oxygen, whilst being kept warm. After observed movement, the mouse was transferred to a heating box (Thermacage, datesand group, UK) prewarmed to 30 °C until normal mouse activity resumed (eating, walking, grooming) when the mouse was returned to group IVC housing.

#### 2.15.4 Micro-CT procedure and analysis of results

Micro-CT was performed using a MILabs OI-CTUHXR preclinical imaging scanner (Utrecht, The Netherlands). In the first study, µCT imaging was performed *in vivo* at weeks 2, 4, 6 and 8 and imaging of excised samples *ex vivo* at week 8. In the study with delayed use of the Laponite/BMP-2 coated scaffolds, scanning was performed *in vivo* at weeks 1, 2, 3, 6, 7 and 8 and imaging of excised samples *ex vivo* at week 8 (settings detailed in **Supplementary Information Table 4**). This method is described in Marshall *et al.* [17]. Briefly, general anaesthesia was induced with 5% isoflurane (VetTech) in an induction box, the mouse was then moved to the imaging bed and maintained at 1.5-2.5% isoflurane, 0.8-1.0 L/min 100% O_2_ throughout imaging. Lubrithal in both eyes prevented eye desiccation. BioVet software monitored respiratory activity (minimum 60 breaths per minute) and the temperature of the scanning bed was set to 34 °C. Mice were imaged using 3 bed positions to µCT scan from the neck to the base of the tail, with a total scan time of approximately 15 minutes. Mice were recovered on the warm imaging bed followed by the heating box. After 8 weeks, to obtain higher resolution images; scaffolds were retrieved from the mice post-mortem and scanned in a specialised sample holder. A density phantom, as a reference for quantification of bone density, was scanned using the same parameters. The μCT reconstructions were obtained via MILabs software (MILabs-Recon v. 11.00). Formation of bone was assessed using Imalytics Preclinical software v3.0 (Gremse-IT GmbH). A gauss filter of 0.5 and bone density threshold for analysis was set equal to the average result from the lower density bone phantom. Data from replicates were analysed using GraphPad Prism 9, version 9.2.0. P values <0.05 were considered significant. All data presented as mean and standard deviation (S.D.).

### 2.16 Histological analysis of CAM and mouse subcutaneous study samples

#### 2.16.1 Processing, embedding and sectioning of samples

Samples were fixed in 4% PFA for 1 day for CAM samples and at least 3 days for mouse subcutaneous samples. Samples from 2 mice were decalcified prior to processing for 5 days with 5% ethylenediaminetetraacetic acid (EDTA) in 0.1 M Tris in dH_2_O (pH 7.3 with NaOH). The sample was taken out of 4% PFA, rinsed in PBS and placed in decalcification solution at 4 °C on the rotator and the solution changed weekly with µCT/x-ray used to confirm complete decalcification prior to rinsing in PBS and further processing. The remaining mice samples were not decalcified to allow Alizarin red and Von Kossa staining and were stored in 70% ethanol prior to processing. After dehydration through ethanol solutions (70%, 90%, 100% twice, with 30 minutes in each concentration) and Histoclear:100% ethanol mix in a 1:1 ratio, followed by 100% Histoclear twice, for 45 minutes in each solution, the samples were processed in a Heraeus vacutherm vacuum oven in molten paraffin wax (Histowax, Leica) at 65 °C for 1 hour. Fresh wax was added and a further 1 hour in the vacuum oven commenced. The samples were embedded in fresh molten wax and cooled at 0-1 °C for 2 hours before storage at 4 °C. The blocks were sectioned at 7 µm on a Microm 330 microtome (Optec, UK). The sections were transferred via a water bath to pre-heated glass slides for 2 hours until dry and placed in the slide oven at 37 °C for 4-6 hours prior to storage at 4 °C.

#### 2.16.2 Histology staining

##### 2.16.2.1 Alcian blue/Sirius red

The staining method has been previously described [17]. The tissue sections on slides were rehydrated through Histoclear (2 x 7 minutes), ethanol solutions of 100% (twice) to 90% to 50% (2 minutes each), followed by immersion in water. Weigert’s Haematoxylin was applied for 10 minutes, removed by immersion 3 times in acid/alcohol (5% HCl/70% ethanol) followed by washing in the water bath for 5 minutes. Slides were immersed in 0.5% Alcian blue 8GX in 1% acetic acid for 10 minutes, 1% molybdophosphoric acid for 10 minutes, followed by rinsing with water prior to staining with 1% Picrosirius Red (Sirius Red) for 1 hour. Excess stain was rinsed off in water and the slides were dehydrated in increasing concentrations of ethanol of 50%, 90%, 100% twice and Histoclear twice for 30 seconds in each prior to mounting with dibutyl phthalate xylene (DPX) and a glass coverslip and allowed to dry.

##### 2.16.2.2 Goldner’s Trichrome

The staining method has been previously described [17]. In brief, the method followed is identical to Alcian blue/Sirius Red staining to the point of acid/alcohol immersion and washing in water for 5 minutes. Ponceau Acid Fuchsin/Azophloxin (Sigma) was applied for 5 minutes followed by a 15 second wash in 1% acetic acid. Phosphomolybdic acid/Orange G was applied for 20 minutes followed by another 15 second wash in 1% acetic acid. Light Green was applied for 5 minutes followed by the third 15 second wash in 1% acetic acid. The sections on the slides were dehydrated in ethanol 90% and 100% twice and Histoclear twice for 30 seconds each prior to mounting with DPX and a glass coverslip and allowed to dry.

##### 2.16.2.3 Von Kossa staining

The staining method has been previously described [17]. In brief, the slides were rehydrated and incubated with 1% silver nitrate under UV light for 20 minutes. Slides were washed in water and incubated with 2.5% sodium thiosulfate for 8 minutes, washed with water and counterstained with Alcian blue for 1 minute. The slides were washed in water and Van Gieson’s stain was applied for 5 minutes. The slides were blotted, dehydrated using ethanol at 90%, 100% twice for 30 seconds each and Histoclear twice for 30 seconds each. Slides were mounted with DPX and a glass cover slip applied and allowed to dry.

##### 2.16.2.4 Alizarin red/light green staining

The staining method has been previously described [17]. In brief, the slides were placed on a slide rack and stained by application of Alizarin red S (40 mM) stain solution for 2 minutes. The slide was blotted and light green counterstain applied to the slide for 2 minutes. This was blotted and acetone applied for 30 seconds followed by acetone/Histoclear (50:50) for 30 seconds, followed by Histoclear for 30 seconds. Slides were mounted with DPX and a glass cover slip was applied and allowed to dry.

Histological samples were imaged using the Zeiss Axiovert 200 digital imaging system using bright field microscopy with the halogen bulb. Images were captured using the Axiovision 4.2 imaging software.

## 3. Results

### 3.1 PCL-TMA900 cytocompatibility and biocompatibility and the efficacy of the Laponite/BMP-2 coating

#### 3.1.1 HBMSCs remain viable, but less readily attach and proliferate on PCL-TMA900 scaffolds compared to PCL-TMA scaffolds and biocompatibility was confirmed by the CAM assay

The PCL-TMA900 scaffold material was tested for cytocompatibility and biocompatibility prior to evaluation of the Laponite/BMP-2 coating on the PCL-TMA900 material *in vivo*. Cytocompatibility of the PCL-TMA900 with HBMSCs was verified by alamarBlue™ HS, as clinical use in human patients is the target application. The PCL-TMA900 scaffolds supported HBMSC growth over a 14-day period (**Figure 1 A)**, although significantly fewer cells were observed to be attached to the PCL-TMA900 scaffold compared with the PCL-TMA scaffold after 24 hours with the live staining, resulting in fewer cells present at day 14 than on the PCL-TMA scaffolds (**Figure 1 B**). The PCL-TMA900 scaffolds were analysed for biocompatibility and the ability to sustain angiogenesis using the CAM assay. Chick viability variation was related to one unformed chick in each group (**Figure 1 C**) and the PCL-TMA900 scaffolds demonstrated the ability to support angiogenesis, with a similar Chalkley score to the PCL-TMA scaffolds (**Figure 1 D**). The tissue around the PCL-TMA900 scaffolds appeared normal, with areas of what appeared to be resolving haemorrhage around the scaffold where the scaffold was embedded in the CAM tissue (**Figure 1 E**). Histologically, the PCL-TMA900 scaffold ridges facilitated interdigitation with the CAM tissue. Mucin in the tissue around the periphery of the PCL-TMA900 scaffolds was observed following Alcian Blue staining while Goldner’s trichrome staining highlighted the vascularity of the CAM surrounding the PCL-TMA900 scaffolds with multiple blood vessels and red blood cells seen with normal collagen tissue architecture within the CAM (**Figure 1 F**).

**Figure 1:**
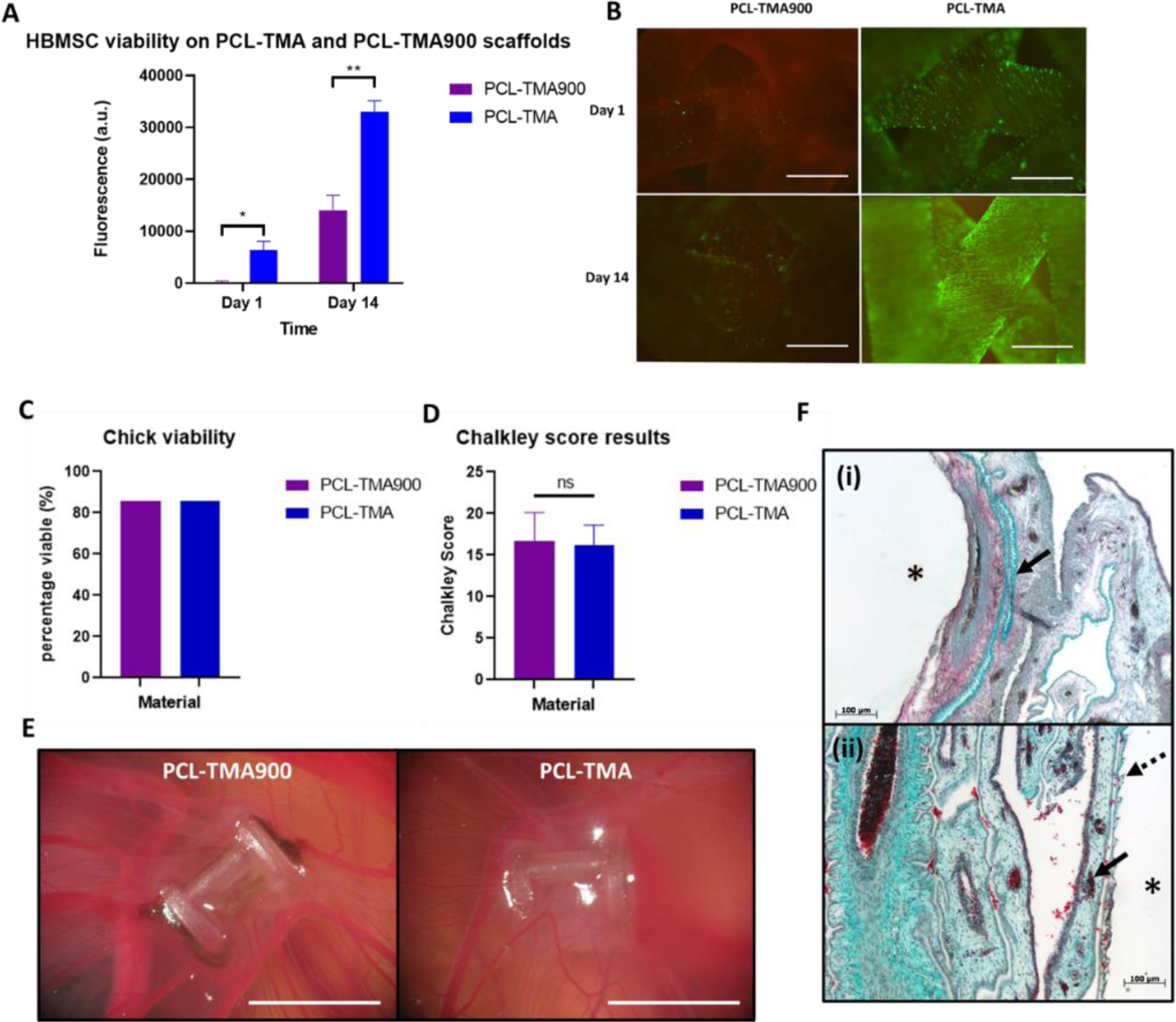
Cytocompatibility and biocompatibility of PCL-TMA900 scaffold material. (A) alamarBlue™ HS fluorescence results of PCL-TMA compared to PCL-TMA900 scaffolds with significantly greater cell metabolism results on the PCL-TMA material (n = 3). Multiple unpaired t-test with Welch correction, mean and S.D. shown, *p<0.05, **p<0.01. (B) The PCL-TMA900 scaffolds had significantly fewer live (green) cells attached at day 1 compared to the PCL-TMA, however the HBMSCs proliferated. The PCL-TMA had significantly greater cell numbers after 14 days of culture evidenced by the fluorescent green cells covering the PCL-TMA scaffold material. Few dead (red) cells were seen. Scale bar 1 mm. (C) CAM assay chick viability was excellent for the PCL-TMA900, comparable to the PCL-TMA scaffold material (n = 7). (D) No significant difference was seen between the Chalkley score between PCL-TMA900 and the PCL-TMA (n = 6), ns=non-significant. A two-tailed unpaired t-test was used for statistical analysis. (E) Photographs of PCL-TMA900 scaffolds and PCL-TMA scaffolds on the CAM. PCL-TMA900 scaffolds had some dark areas of possible resolving haemorrhage from scaffold implantation and both PCL-TMA materials illustrated biocompatibility and support for angiogenesis, with the scaffolds fully surrounded and integrated with the CAM tissue. Scale bar 4 mm. (F) (i) Alcian blue and Sirius red staining of PCL-TMA900 scaffold (*) with blue mucin (arrow) within the red collagenous matrix. Scale bar 100 µm. (ii) Goldner’s trichrome staining of PCL-TMA900 scaffold (*) with surrounding blood vessels (arrow) and interdigitating tissue with the scaffold surface shape (dashed arrow). Scale bar 100 µm.

#### 3.1.2 Laponite persists on the PCL-TMA900 material and facilitates protein binding to the PCL-TMA900 scaffold

To determine if the Laponite remained adherent to the scaffold over time, ethidium homodimer staining of the Laponite was performed. The Laponite remained present on the scaffold surface for up to 7 days of incubation in PBS as detected using red ethidium homodimer stain on/within the Laponite (**Figure 2 A**). The uncoated control scaffolds showed no ethidium homodimer staining at any time examined. The Laponite coated scaffolds in PBS for 1 hour simulated the minimum time the scaffolds were expected to be immersed in BMP-2/PBS when coating the scaffolds with the growth factor solution (**Figure 2 A**). Fluorescent FITC-BSA was used as a model protein to determine the distribution of protein after the Laponite coated scaffolds were immersed in a protein-containing solution. The PCL-TMA900 scaffold did not fluoresce in PBS or BSA/PBS; however, FITC fluorescence of the PCL-TMA900 scaffold was observed when immersed in the FITC-BSA solution. The Laponite coated PCL-TMA900 scaffold displayed bright, focal fluorescent areas over the entire PCL-TMA900 scaffold, indicating the fluorescent protein was concentrated in these regions by the concurrent Laponite coating (**Figure 2 B**). To assess the amount of BMP-2 binding to uncoated scaffolds and Laponite coated scaffolds, an ELISA was performed to measure residual BMP-2 from 5 µg/mL or 1 µg/mL solutions that were used to coat the scaffolds for the ALP staining study to evaluate C2C12 bioactivity. The results confirmed BMP-2 was sequestered from the solution, with less BMP-2 remaining in the solution when Laponite coated scaffolds were immersed, compared to when uncoated scaffolds were immersed. This effect was observed at both 5 µg/mL and 1 µg/mL concentrations of BMP-2 with a similar percentage of BMP-2 taken up by the Laponite (77.0 ± 0.3% and 76.0 ± 4.1% respectively) (**Figure 2 C**).

**Figure 2:**
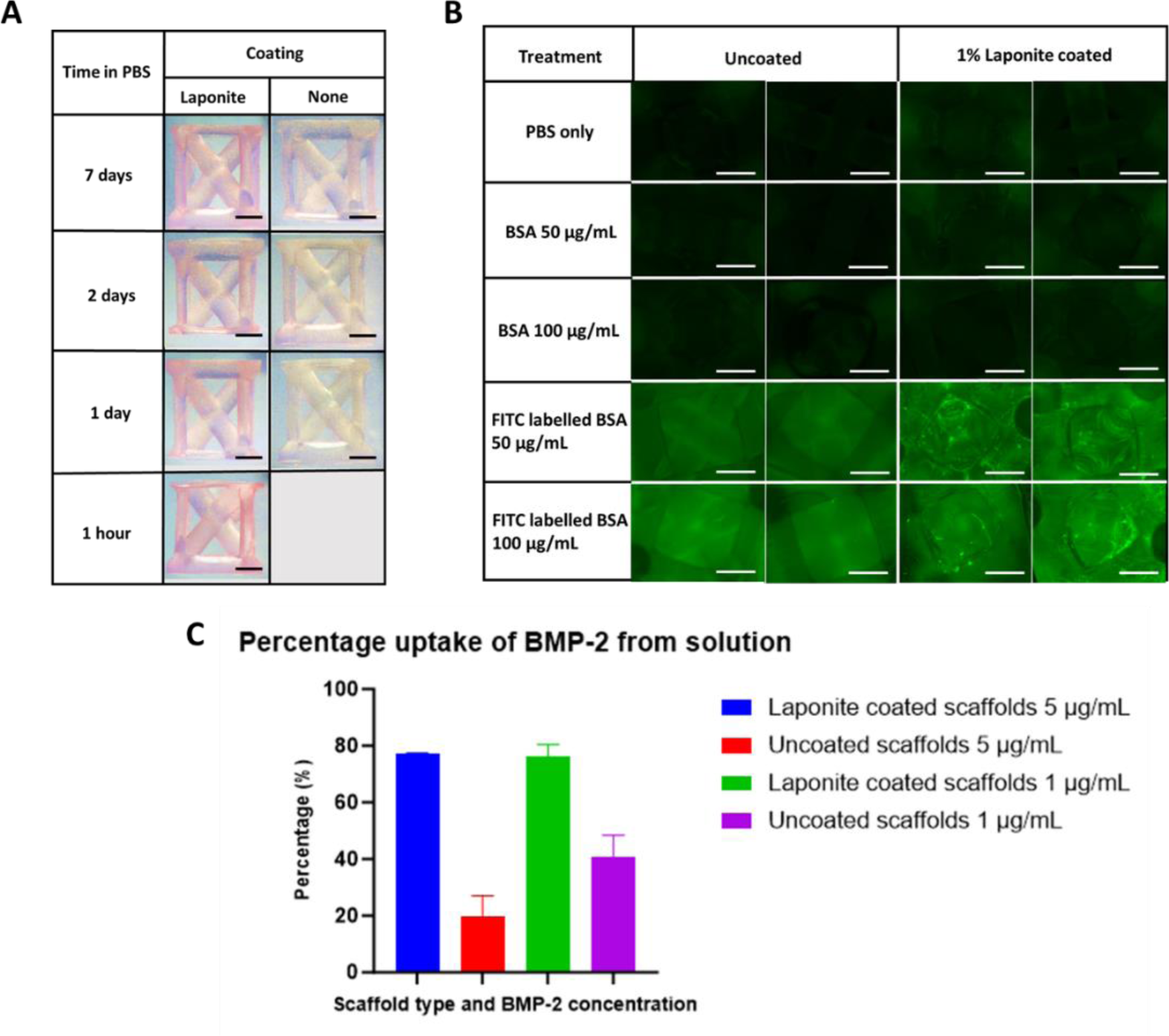
Confirmation of Laponite persistence and protein uptake on PCL-TMA900 scaffolds. (A) Ethidium homodimer staining confirmed the persistence of Laponite on the PCL-TMA900 scaffold as Laponite coated scaffolds maintained a red colour (left column) when the ethidium homodimer bound to the Laponite, while there was no red colour on the uncoated control scaffolds (right column), images manipulated to saturation 200% for each image to highlight the colour difference, n = 3 for each condition, scale bar 1 mm. (B) Visualisation of FITC-BSA uptake in regions of Laponite coated scaffolds. The PCL-TMA900 scaffold itself did not show autofluorescence allowing visualisation of the FITC-BSA protein, which appeared as a green background colour. However, PCL-TMA900 scaffolds with the Laponite coating showed a bright fluorescent green FITC-BSA signal attached to the scaffold indicating the Laponite/FITC-BSA combination on the surface, n = 2, scale bar 1 mm. (C) The percentage uptake of BMP-2 from the coating solutions. The Laponite coated scaffolds were able to sequester more BMP-2 from the solution than the uncoated scaffolds at 1 µg/mL and 5 µg/mL BMP-2 concentrations. Uncoated and Laponite coated scaffold samples were plated out once each as biological triplicates, mean and S.D. shown, n = 3.

#### 3.1.3 Laponite on PCL-TMA900 scaffolds sequester active BMP-2, inducing osteogenic differentiation of C2C12 cells, while delayed use of the Laponite/BMP-2 coating reduced bioactivity in vitro

BMP-2 dose dependent differentiation of murine myoblast C2C12 cells, as evidenced by red ALP staining, was assessed to ensure the C2C12 cells were capable of osteogenic differentiation (**Supplementary information Figure 7**). C2C12 cells were selected due to their widespread use as a marker cell type for confirming the activity of BMP-2. The ability for Laponite to bind to the BMP-2 on a 3D scaffold and retain bioactivity was assessed compared to when uncoated scaffolds were immersed in BMP-2, producing BMP-2 only coated scaffolds. Concurrently, the effective concentration of BMP-2 (5 µg/mL or 1 µg/mL) was assessed following immersion of PCL-TMA900 scaffolds for 24 hours in BMP-2 solution. This determined if a less concentrated BMP-2 solution could produce a similar positive result, reducing the mass of BMP-2 required if the scaffold was saturated at a lower concentration/mass of BMP-2. In the ALP staining assay using the 3D scaffolds, the C2C12 cells on the Laponite coated PCL-TMA900 scaffolds displayed marked ALP expression in response to the bound BMP-2 when 5 µg/mL of BMP-2 was used. There was noticeably less ALP expression when PCL-TMA900 Laponite coated scaffolds were coated using 1 µg/mL of BMP-2 and no ALP expression was evident when there was no BMP-2 present. The BMP-2 only control scaffolds showed very low ALP expression when 5 µg/mL BMP-2 was used and negligible ALP expression in response to the bound BMP-2 when 1 µg/mL of BMP-2 or no BMP-2 were used for scaffold immersion. The positive control scaffolds exposed to 5 µg of BMP-2 in the media displayed more marked ALP expression in the Laponite group than in the BMP-2 only group as the Laponite sequestered the BMP-2 to the surface of the scaffold during the culture period (**Figure 3 A**). To confirm that the ALP staining was not absent due to a lack of cell confluence on the BMP-2 only scaffolds, ethidium homodimer staining confirmed C2C12 cells covering the scaffold surface, imaged by fluorescence microscopy (**Figure 3 A**). To investigate if the dry Laponite/BMP-2 coating was stable for a prolonged period, to explore possible ready-to-use clinical applications of the coating, the Laponite coating was applied followed by the BMP-2 coating, with the well plate stored at RT for 24 hours. When examined and assessed after 24 hours, there was no appreciable ALP staining compared to cells seeded immediately post 2 hour drying time. This may be due to loss of activity of the BMP-2 upon the Laponite surface, with only BMP-2 within the Laponite present which would not be accessible to the C2C12 cells without digestion of the Laponite and release of the BMP-2 by macrophages, hence going forward in this work to recapitulate this experiment *in vivo*. The positive control wells showed marked positive red ALP stain, which was more pronounced in the centre of the well with a lighter staining ring at the edge of the well. We hypothesise this was a result of the increased Laponite meniscus during the coating process, therefore the Laponite took up the BMP-2 from the media, limiting BMP-2 access by the C2C12 cells as the cells did not penetrate the Laponite *in vitro* (**Figure 3 B**).

**Figure 3:**
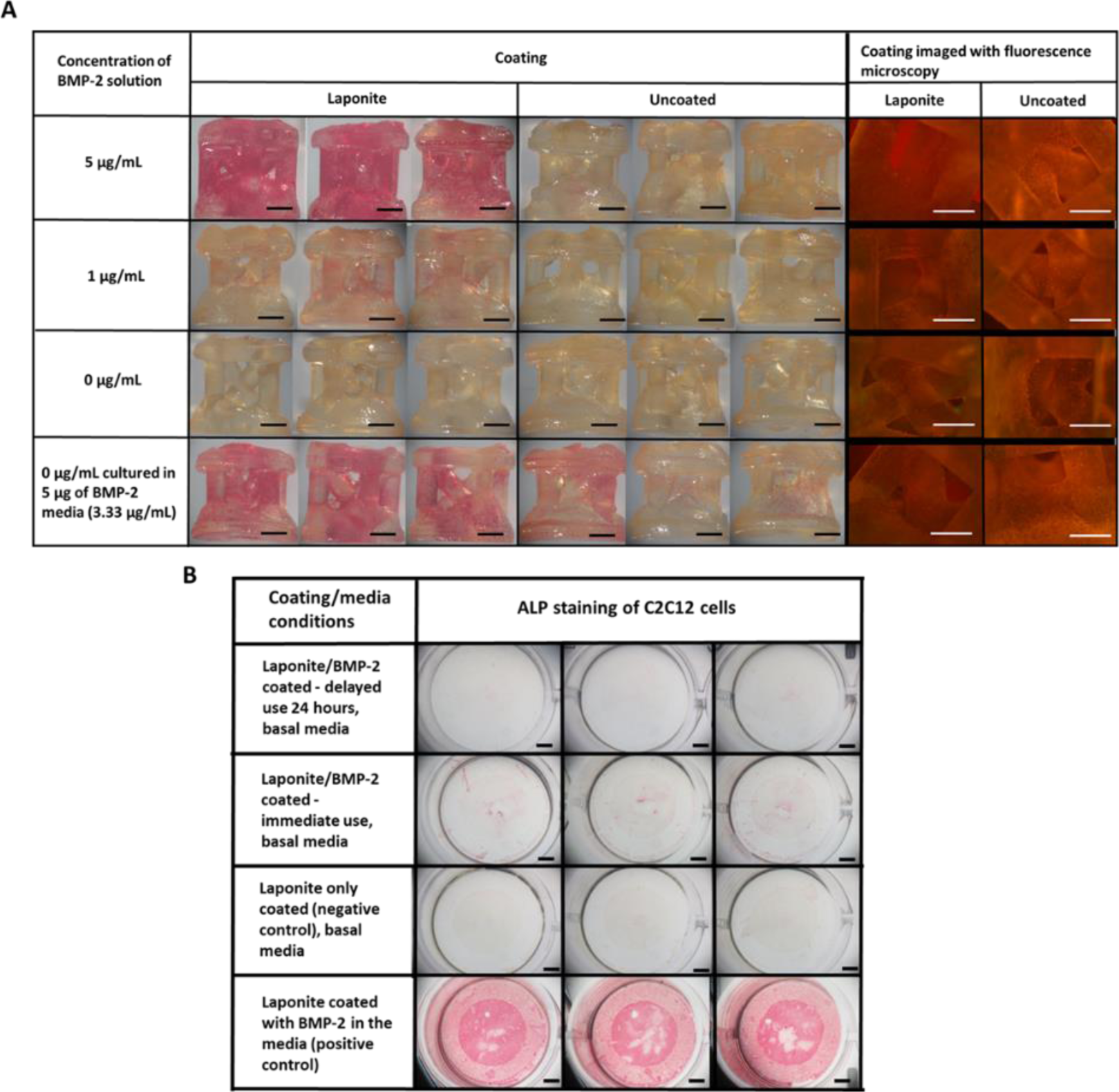
Confirmation of BMP-2 binding and bioactivity on Laponite coated PCL-TMA900 scaffolds and results of coating use after 24 hours. (A) Photographs of C2C12 ALP and ethidium homodimer staining on Laponite coated and uncoated PCL-TMA900 scaffolds. Positive ALP expression results displayed by the vivid red colour on the Laponite coated scaffolds immersed in 5 µg/mL BMP-2, with less ALP expression from C2C12 cells when 1 µg/mL was used. The negative controls without BMP-2 showed no ALP expression. The uncoated scaffolds displayed low, inconsistent ALP expression. The positive controls showed ALP expression, with greater clarity and consistency in the Laponite coated group compared to the uncoated control group, n = 3, scale bar 1 mm. Fluorescence microscopy using ethidium homodimer staining of C2C12 cell nuclei on PCL-TMA900 scaffolds, post ALP staining, displayed fixed cells upon all scaffolds in all groups by bright red dots of the nucleus stain. Representative images shown for the fluorescence microscopy images, n = 3. Scale bar 1 mm. (B) Bioactivity of BMP-2 after 24 hours of drying time on Laponite coated wells. The BMP-2 appeared to have lost bioactivity after remaining dry at room temperature for 24 hours prior to cell seeding, evidenced by lack of ALP staining compared to those wells in which BMP-2 was dried for 2 hours prior to immediate cell seeding. The positive control wells with free BMP-2 in the media showed a marked increase in red ALP staining whereas the negative control wells demonstrated no red ALP staining (n = 3, scale bar 1 mm).

### 3.2 Subcutaneous bone formation in the Laponite/BMP-2 coated PCL-TMA900 scaffolds compared to control and BMP-2 only coated scaffolds

#### 3.2.1 Bioactive BMP-2 is sequestered by Laponite on PCL-TMA900 scaffolds producing bone around the subcutaneous Laponite/BMP-2 coated PCL-TMA900 scaffolds

The subcutaneous implantation study in mice demonstrated that the uncoated and BMP-2 only coated PCL-TMA900 scaffolds displayed no detectable mineralisation at any time point examined. In comparison, bone formation was observed around all the Laponite/BMP-2 coated scaffolds as well as the positive control collagen sponge soaked with 5 µg BMP-2 from week 2 onwards (**Figure 4 A**). The response to the collagen sponge/BMP-2 and Laponite/BMP-2 coated scaffolds was quantified using micro CT. Mouse 3 had a marked rapid response to the BMP-2 within the collagen sponge from week 2, with remodelling over time, while the other 3 mice showed a gradual increase in bone volume from week 2 to week 8 (**Figure 4 B**). Mouse 2 displayed a rapid and marked response to the BMP-2 on the Laponite coated scaffold with no obvious change in bone formation between weeks 2 and 4, followed by a further increase in mineralisation between weeks 4 and 6 with a smaller increase between weeks 6 and 8. The other 3 mice displayed a gradual increase in bone formed from week 2 to 8 (**Figure 4 C**). The materials scanned *ex vivo* showed no significant bone formation in response to either of the uncoated PCL-TMA900 scaffolds or BMP-2 only coated PCL-TMA900 scaffolds. However, the Laponite/BMP-2 coated PCL-TMA900 scaffolds and the collagen sponge with BMP-2 displayed significant bone formation, in agreement with findings when the constructs were scanned at lower resolution *in vivo* (**Figure 4 D**). A shell of bone surrounding the Laponite/BMP-2 coated scaffold could be clearly seen and the collagen sponge displayed bone forming an outer shell (**Figure 4 E**). The BMP-2/PBS coating solution remaining from the scaffold coating process was examined to determine the mass of BMP-2 delivered by the scaffolds. The ELISA results showed the Laponite coated scaffolds took up a far greater percentage of BMP-2 than the BMP-2 only coated scaffolds from the starting stock solution of 5 µg/mL which corresponded with the degree of bone formation on each scaffold coating (**Figure 4 F**).

**Figure 4:**
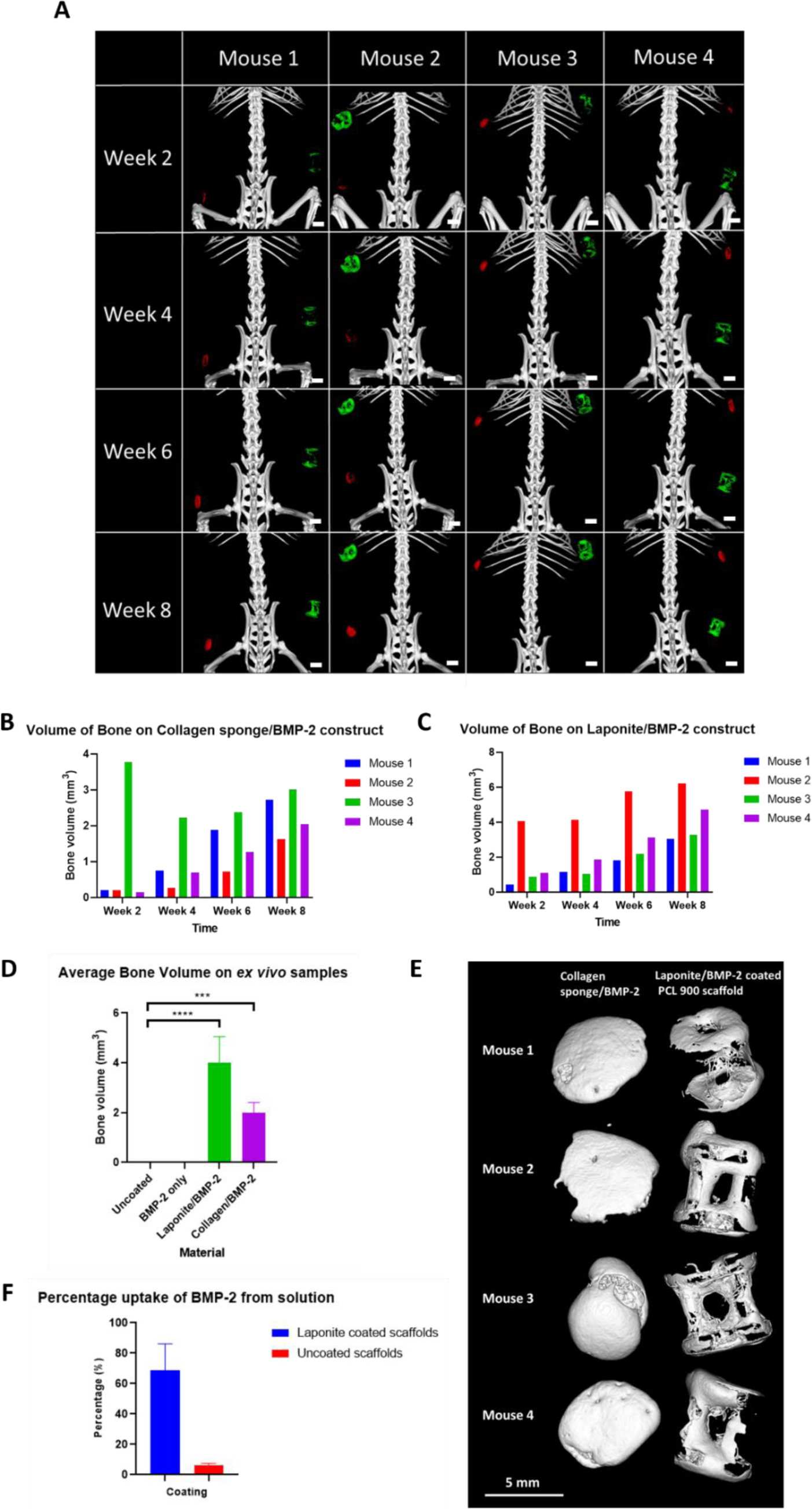
Murine subcutaneous model examining Laponite/BMP-2 coated PCL-TMA900 scaffolds. (A) µCT images display the collagen sponge (red) from week 2 onwards with a progressive increase in volume of mineralisation over time, while the Laponite/BMP-2 coated scaffolds (green) were seen as a faint shell in mice 1, 3 and 4, with more extensive mineralisation observed in mouse 2 at week 2 and this continued to progress in each mouse over the remaining 6-week study period (n = 4), scale bar 5 mm. (B) Volume of bone formed on collagen sponge/BMP-2 scaffolds. Mouse 3 displayed a rapid response to the BMP-2 in the collagen sponge with remodelling of the volume of bone over time, while the other mice displayed a gradual increase in mineralisation of the collagen sponge from week 2 to week 8 (n = 4). (C) Volume of bone formed on Laponite/BMP-2 coated scaffolds. Mouse 2 displayed a rapid marked response to the BMP-2 held on the laponite coated scaffold, with increasing bone volume between week 4 to week 8, while the other 3 mice displayed a gradual increase in bone formation around the scaffold between week 2 and week 8 (n = 4). (D) Average bone formation on *ex vivo* constructs. Compared to uncoated PCL-TMA900 scaffolds, the Laponite/BMP-2 and collagen sponge/BMP-2 constructs had significantly greater bone while the BMP-2 only control displayed negligible bone formation (n = 4). One-way ANOVA with Dunnett’s multiple comparisons test, mean and S.D. shown. (E) *Ex vivo* µCT images of collagen sponge/BMP-2 and Laponite/BMP-2 coated scaffolds at week 8 timepoint. The collagen sponge/BMP-2 mineralised at the periphery with an extra protrusion of bone in Mouse 3. The Laponite/BMP-2 showed a shell of mineralisation which was not completely uniform around the scaffold, but most of the scaffold appeared to be coated (n = 4), scale bar 5 mm. (F) ELISA results of BMP-2 uptake by Laponite coated and dH_2_O immersed (uncoated) PCL 900 scaffolds. The percentage of BMP-2 uptake was greater in the Laponite coated scaffold group, n = 4 biological replicates for each scaffold type with each sample plated out in triplicate wells and the average result of each scaffold triplicate is shown, mean and S.D. shown.

Histological analysis confirmed the µCT results, with no mineralisation found around the uncoated PCL-TMA900 scaffolds or BMP-2 only coated PCL-TMA900 scaffolds. Constructs which were not demineralised showed a black mineralised shell on the Laponite/BMP-2 coated scaffolds upon Von Kossa staining and red mineral upon Alizarin Red staining, consistent with findings on the positive control collagen sponge/BMP-2 (**Figure 5**). Decalcified collagen sponge/BMP-2 and Laponite/BMP-2 coated scaffold samples showed a rim of bone tissue with regularly distributed osteocytes seen on Alcian blue/Sirius red staining and bright red osteoid was seen on Goldner’s Trichrome staining (**Figure 5**).

**Figure 5:**
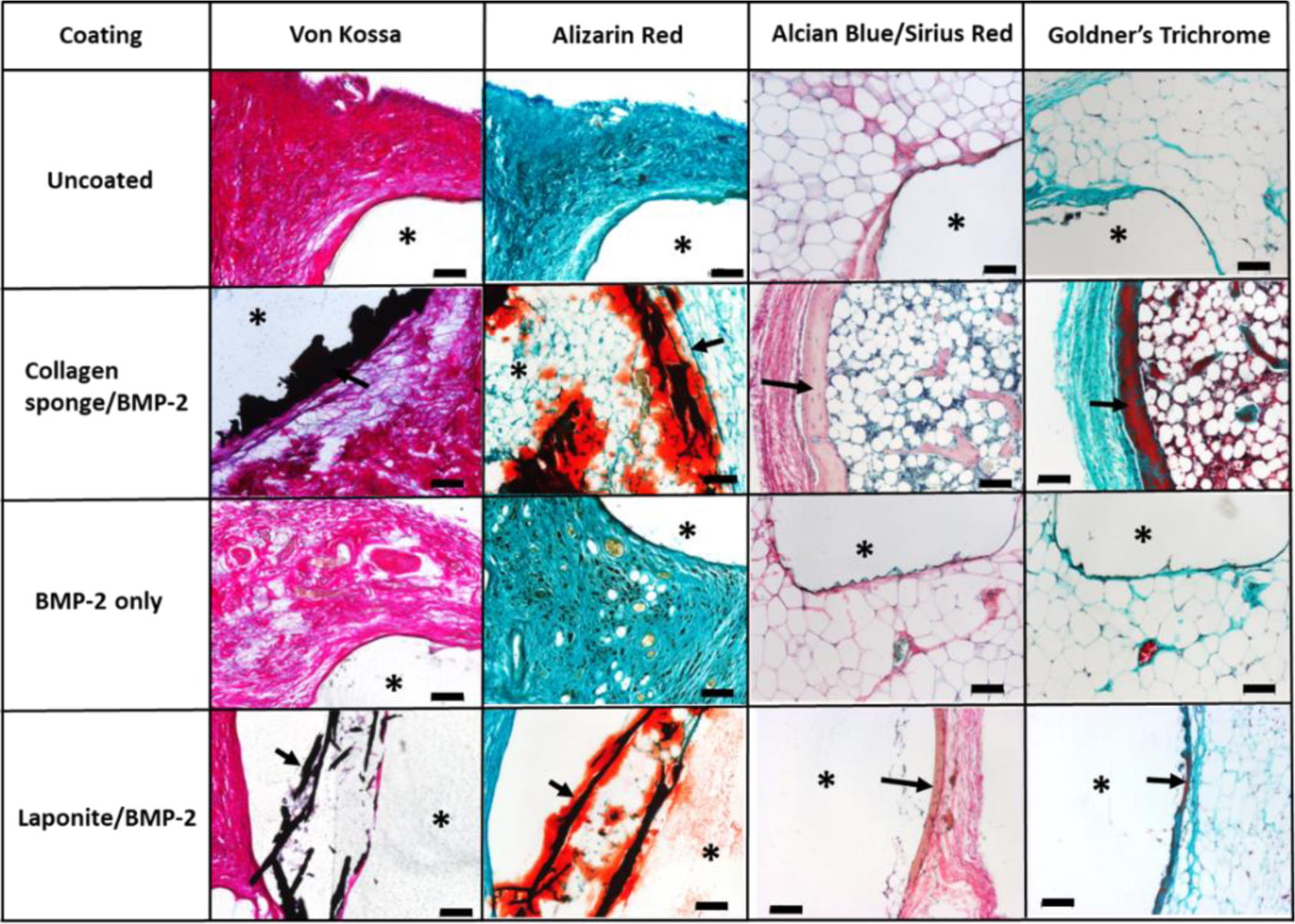
Histology results of the subcutaneous Laponite/BMP-2 coated PCL-TMA900 scaffold implantation study. The black (Von Kossa) and red (Alizarin red) staining of bone (arrows) was seen in the collagen sponge/BMP-2 and Laponite/BMP-2 groups only. Alcian Blue and Sirius red staining with osteocytes and Goldner’s trichrome showing osteoid (arrows) in the collagen sponge/BMP-2 and Laponite/BMP-2 groups only. (*) denotes the PCL 900 scaffold/collagen sponge. Scale bar 100 µm.

#### 3.2.2 Delayed implantation of Laponite/BMP-2 coated PCL-TMA900 scaffolds produced reduced bone volume and less rapid bone formation, than the immediate use of Laponite/BMP-2 coated scaffolds

This mouse subcutaneous implantation study indicated the potential of using the Laponite/BMP-2 coated scaffolds after 24 hours, however, the response was not as marked or rapid as when the scaffolds were used immediately once dry post coating. The negative control uncoated PCL-TMA900 scaffolds contained no detectable mineralisation at any time point on µCT analysis *in vivo* or *ex vivo*, however, the positive control of collagen sponge soaked with 5 µg BMP-2 mineralised in all 4 mice by week 8. The Laponite/BMP-2 coated scaffolds used immediately showed marked mineralisation, with less obvious bone formation in the delayed Laponite/BMP-2 coated group when scanned *in vivo* (**Figure 6 A**). For the collagen sponge/BMP-2 implants, mice 2 and 3 were the only mice to show a mineralisation response at week 2, while in the previous study all mice showed a response at this timepoint. This difference may be due to the BMP-2 being kept in buffer solution for a prolonged period. However, by week 3 all mice displayed mineralisation of the collagen sponge occurring which continued as a gradual sustained increase in mice 2 and 4, while mice 1 and 3 had a marked increase in mineral volume between weeks 3 and 6 followed by a reduction, with a large increase between weeks 7 and 8 (**Figure 6 B**). In the Laponite/BMP-2 coated scaffolds used immediately once dry, mice 2 and 3 displayed a rapid marked response to the BMP-2 on the Laponite coated scaffold with minimal change between weeks 2 and 3, followed by a further increase in mineralisation between weeks 3 and 6 which became more stable over the following 2 weeks until week 8. Mice 1 and 4 had a minimal response until week 6, with similar results at week 7 followed by an increase in bone volume at week 8 (**Figure 6 C**).

**Figure 6:**
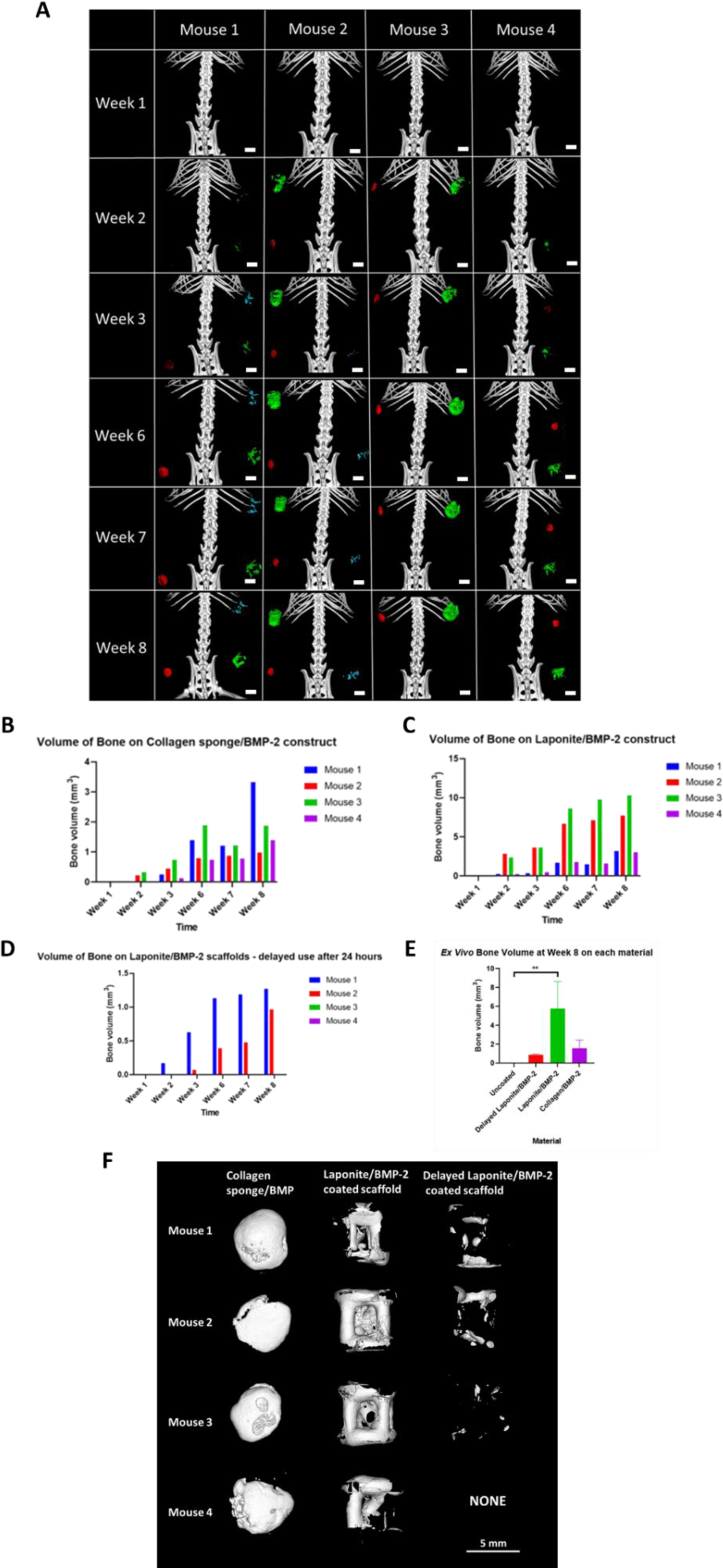
µCT image results of Laponite/BMP-2 and delayed use Laponite/BMP-2 coated PCL-TMA900 scaffolds in the murine subcutaneous model. (A) The collagen sponge could be seen in red in all mice from week 3 onwards with a progressive increase in volume of mineralisation over time, while the Laponite/BMP-2 coated scaffolds could be seen in green as a faint shell in all mice from week 2 onwards, n=4 each group. The delayed use of Laponite/BMP-2 coated scaffolds shown in blue, gave interesting results of a slower response time and less bone volume, but a response materialised over time in mice 1 and 2 *in vivo* (n = 3, 24 hours delayed use, n = 1, 6 months delayed use in mouse 4), scale bar 5 mm. (B) Volume of bone formed on collagen sponge/BMP-2 scaffold. Mouse 1 displayed a slow response to the BMP-2 in the collagen sponge with remodelling of the volume of bone over time, followed by a rapid massive response between weeks 7 and 8, while mice 2 and 4 had a stepwise increase in mineralisation of the collagen sponge from week 2 to week 8. Mouse 3 showed fluctuations in bone volume over time, but the sponge did mineralise in all mice during the 8-week study period (n = 4). (C) Volume of bone formed on Laponite/BMP-2 coated scaffold. Mice 2 and 3 had a rapid marked response to the BMP-2 held on the laponite coated scaffold, compared to mice 1 and 4 who had less mineralisation formed on the scaffold over time (n = 4). (D) Volume of bone formed on Laponite/BMP-2 coated scaffolds after delayed implantation. Only mice 1 and 2 showed bone formation on the scaffolds maintained dry for 24 hours prior to implantation which formed slowly followed by a large increase between weeks 3 and 6 in both mice, with a further large increase in mineralisation in mouse 2 by week 8 (n = 3 implanted after 24 hours, n = 1 implanted after 6 months. (E) Average volume of bone on *ex vivo* constructs. Compared to uncoated PCL-TMA900 scaffolds, the Laponite/BMP-2 had significantly greater bone while the Delayed Laponite/BMP-2 group had no significant difference in bone formation, n = 3, One-way ANOVA with Dunnett’s multiple comparisons test, **p<0.01, mean and S.D shown. (F) *Ex vivo* µCT images of collagen sponge/BMP-2, Laponite/BMP-2 and Delayed Laponite/BMP-2 coated scaffolds at the week 8 timepoint. The collagen sponge/BMP-2 displayed a smooth shell of bone in a disc shape with an extra protrusion of bone in mouse 4. The Laponite/BMP-2 formed a shell of bone as per the previous study, while the Delayed Laponite/BMP-2 group had discrete patches of bone forming on the scaffold (n = 4), scale bar 5 mm.

In contrast, the Laponite/BMP-2 coated scaffolds used in a delayed manner, 24 hours post coating, showed a mineralisation response in 3 of the 4 mice, with mineralisation volumes less than when the coating was used immediately. Bone was observed to take 8 weeks to form in mouse 3, showing a delayed response to the BMP-2 on the scaffold and could only be detected upon higher resolution µCT image reconstruction. In the Laponite/BMP-2 delayed use scaffold group, mouse 1 was the first to respond to the BMP-2 on the scaffold with a marked increase in bone volume until week 6, while mouse 2 began to show mineralisation at week 3 followed by an increase to week 6 and a further large increase to week 8. Mice 3 and 4 displayed no detectable bone formation on these delayed use PCL-TMA900 scaffolds upon µCT scanning *in vivo* (**Figure 6 D**).

The samples imaged *ex vivo* had no mineralisation in response to the uncoated PCL-TMA900 scaffolds. The results of average bone volume for mice 1-3 showed a significant difference in bone volume formed on Laponite/BMP-2 coated scaffolds compared to the uncoated PCL-TMA900 scaffolds. The collagen sponge/BMP-2 constructs did not show a significant difference in this study. Mouse 4 was omitted from this data analysis due to the scaffold used for delayed implantation differing from the other 3 mice (*6 months stored dry versus 24 hours*), however, regardless no bone formed on this delayed use Laponite/BMP-2 coated scaffold (**Figure 6 E**). A shell of bone surrounding the Laponite/BMP-2 coated scaffold could be clearly seen and the collagen sponge had a mineralised outer shell in all mice, while the delayed Laponite/BMP-2 coated scaffolds showed mineralisation preferentially along the cylindrical ‘ends’ of the scaffolds at the periphery (**Figure 6 F**).

Histological analysis confirmed the findings of the µCT images with a mineralised shell confirmed around the Laponite/BMP-2 coated scaffolds, Delayed Laponite/BMP-2 coated scaffolds and positive control collagen sponge/BMP-2 upon Von Kossa and Alizarin Red staining (**Figure 7**). Decalcified samples of the collagen sponge/BMP-2, Laponite/BMP-2 and Delayed Laponite/BMP-2 coated scaffolds displayed a rim of bone with osteocytes on Alcian blue/Sirius red staining and bright red osteoid seen on Goldner’s Trichrome staining, compared to no bone found on the uncoated PCL-TMA900 scaffold (**Figure 7**).

**Figure 7:**
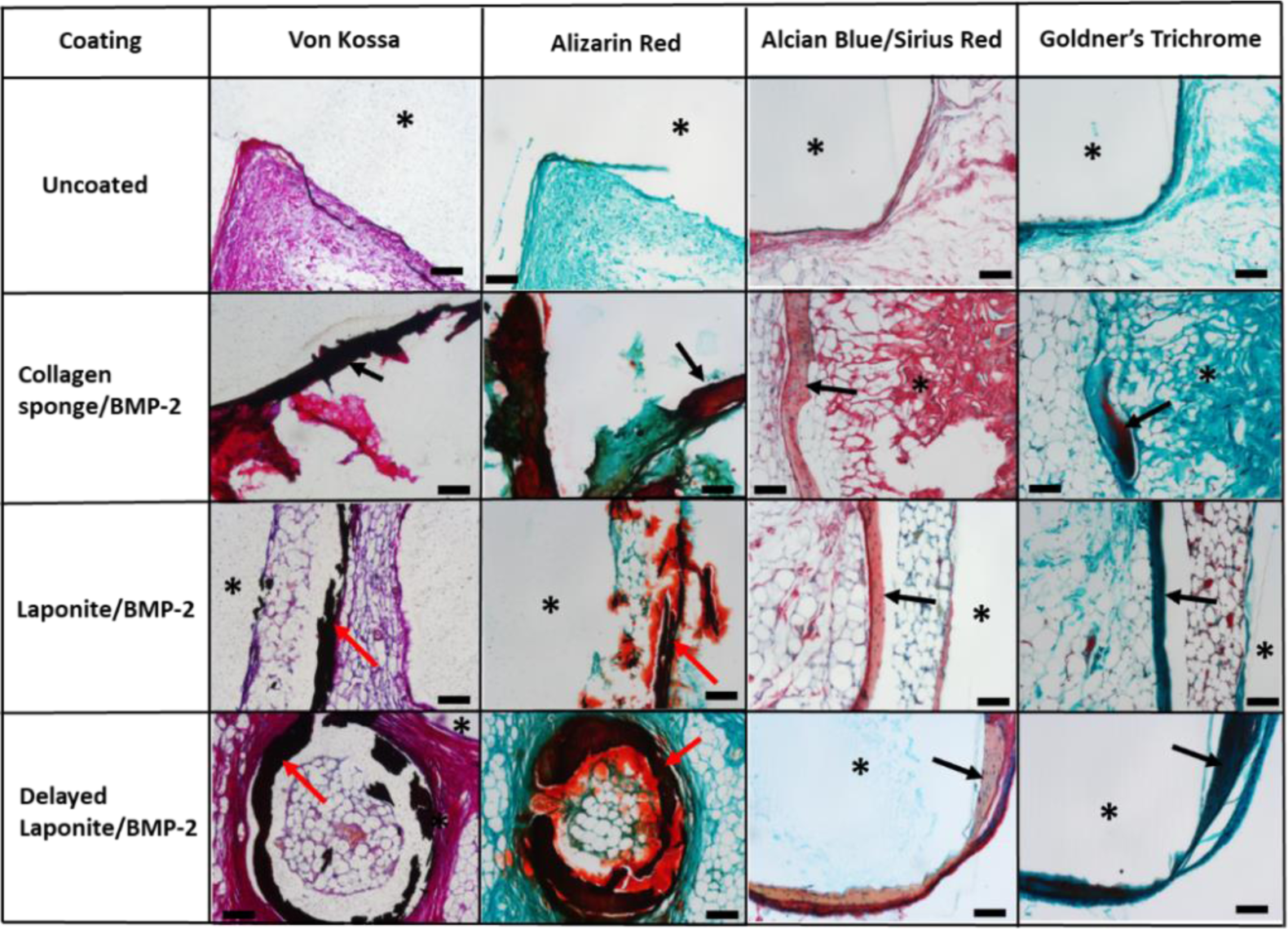
Histology images of bone formation in the murine subcutaneous implantation study. The Von Kossa staining and Alizarin Red staining show black and red staining of bone respectively in the collagen sponge/BMP-2, Laponite/BMP-2 and Delayed Laponite/BMP-2 groups (red arrows), with no staining in the uncoated PCL-TMA900 group. The Alcian Blue and Sirius red staining and Goldners trichrome staining with osteocytes and osteoid staining of bone respectively around the periphery of the collagen sponge/BMP-2, Laponite/BMP-2 groups and Delayed Laponite/BMP-2 groups (black arrows). (*) denotes the scaffold/collagen sponge, scale bar 100 µm.

## 4. Discussion

In the current study, we hypothesised that Laponite nanoclay would adhere to the PCL-TMA900 material and subsequent immersion of the Laponite coated scaffolds in BMP-2 solution would enable sequestration of the BMP-2 by the Laponite coating. This would subsequently produce a bioactive, osteoinductive scaffold, a result confirmed in this study. The Laponite was readily applied and adhered to the PCL-TMA900 scaffolds and was able to bind and deliver active BMP-2 to C2C12 cells, to induce cellular differentiation. The use of Laponite as a dry film has already been demonstrated by Gibbs *et al.*, however Laponite sequestered BMP-2 from surrounding media which, induced osteogenic differentiation of C2C12 cells [18]. The coating method used in this current work dried the BMP-2 on the surface of the Laponite and therefore the BMP-2 was held on the scaffold surface without reliance on addition of BMP-2 solutions at point of use, which could lead to less precise bone formation in the desired *in vivo* location. The *in vitro* results demonstrated that Laponite with BMP-2 had the potential to be a successful coating on the PCL-TMA900 material in the *in vivo* models.

The BMP-2 produced varying results of ALP staining depending on the concentration of BMP-2 and duration of scaffold immersion. Immersion for 24 hours using 5 µg/mL was shown to be sufficient, as assessed by the ALP staining response, therefore, the binding of BMP-2 protein to the clay nanoparticles was relatively rapid [26]. The 24 hour incubation time was practical for experiment preparation when translating from *in vitro* to *in vivo* work. The BMP-2 only coated scaffolds exhibited discrete ALP staining as PCL-TMA900 itself allowed BMP-2 attachment, however, the Laponite coated scaffolds produced the most intense ALP expression, with C2C12 cells visualised by ethidium homodimer staining of each scaffold. A key aim in this study was to demonstrate the complimentary properties of the Laponite and BMP-2 together, with Laponite aiding cell adhesion and retention of BMP-2 upon the surface, allowing the BMP-2 to induce osteogenic differentiation of the surrounding cells over time.

The scaffolds were allowed to dry after BMP-2 application and therefore it was hypothesised that if the scaffolds were maintained in suitable conditions, e.g., dry, in a sterile container, coated scaffolds could be used later, providing a practical, packageable scaffold for clinical use in bone tissue engineering. However, the current studies showed the BMP-2 displayed a loss in bioactivity or adherence to the dry Laponite during the 24 hours. While electrostatic interactions are likely the dominant mechanism of securing proteins to the surface, van der Waals forces, hydrophobic interactions and adsorption of the protein onto the clay particles are also important [26]. From the assessment of Laponite persistence on the scaffold material, the ability for the Laponite coated scaffolds to be stored dry would be beneficial as a growth factor coating could be applied later, with a variety of growth factors to choose from e.g., vascular endothelial growth factor (VEGF), BMP-6, fibronectin (FN) etc. depending on clinical application of the Laponite coating and underlying scaffold material.

A uniform coating method with Laponite would be favourable to immersion and static drying, especially on a hydrophobic material such as PCL-TMA900. Laponite has been crosslinked to polyethylene oxide (PEO) to create a coating for polylactic acid (PLA) nanofibers, coated by immersion in the Laponite/PEA solution which improved hydrophilicity of the PLA material [27]. Laponite has also been mixed with cobalt chloride solution and using carboxymethyl chitosan (CMC) to immobilize the Laponite/cobalt chloride mix to the modified PCL nanofibers of the scaffold [28]. Laponite was combined with PCL as a coating to be applied by electrophoresis to stainless steel implants to improve material properties of metallic bone implants and thus a number of options for incorporating Laponite into polymer coatings exist, however, none have been found to use Laponite alone [29]. In future studies, creating a uniform Laponite coating by use of spraying, spin coating, constant rotation of the scaffold while drying to prevent the coating preferentially collecting at the most dependent part of the scaffold, or the use of a vacuum in very porous materials to encourage the Laponite to be drawn within the pores of the material could be investigated. Additionally, the use of critical point drying or freeze drying could be investigated as options to dry the scaffolds more uniformly without damaging the Laponite or BMP-2 coatings.

The BMP-2 was not rinsed off the surface of the scaffolds after coating to allow, potentially, a high concentration of physically adsorbed ‘free’ BMP-2 on the surface. This ‘free’ BMP-2 would be in addition to having bound BMP-2 within the Laponite, as it is rehydrated during the coating process, providing sustained or delayed release of the protein. Therefore, it was hypothesised that there was a burst release of ‘free’ BMP-2 followed by more sustained release, which was observed in the subcutaneous implantation model. However, after 24 hours of scaffold storage, there was an apparent lack of rapid bone formation due to the loss of a ‘burst’ response as the ‘free’ BMP-2 was perhaps not active/present. In contrast, washing of protein coated Laponite has been shown to preserve the activity of the selected bound protein, such as VEGF in media added to Laponite for 2 hours to allow binding of the protein, followed by washing and seeding of HUVECs. The endothelial tubule network formed by the HUVECs was comparable between the bound VEGF and the positive control of free VEGF in media illustrating that an efficacious concentration remained bound to the Laponite surface [30].

The CAM assay was used to assess the biocompatibility of the uncoated PCL-TMA900 scaffold material, critical to verify prior to subsequent *in vivo* evaluation. The PCL-TMA900 scaffolds were found to be biocompatible with no chick viability concerns following incubation with the scaffolds. Histological examination provided a key tool to assess the infiltration of blood vessels into the PCL-TMA900 construct applied on the CAM and to determine the lack of a foreign body reaction. Repetition of the CAM assay with coated PCL-TMA900 scaffolds was not performed as Laponite and BMP-2 are known to be biocompatible and the scaffolds were not testing enhancement of angiogenesis due to the scaffold coating, rather osteogenesis. However, the entire scaffold was not in contact with the CAM and therefore the CAM does not provide a substitute for the mouse studies in assessment of bone formation. Further, in keeping with the application of the ‘3Rs’ there would be no benefit in using more chicks to gain more data, which does not inform or translate to the dose of BMP-2 to apply to the bone defect preclinical animal models [31, 32].

The mouse subcutaneous implantation model allowed scaffold assessment in a rodent preclinical model and confirmed the biocompatibility of the PCL-TMA900 scaffolds and coating. Reporting the concentration and volume of growth factor use in animal models and details of material size and surface area provides a more accurate comparison between animal studies, especially subcutaneous implant studies, as these can vary as previously reviewed by Gothard *et al.* [33].

The initial subcutaneous implantation study found the Laponite/BMP-2 coated PCL-TMA900 scaffolds demonstrated bone formation on all scaffolds when the coating was used immediately, while the controls of BMP-2 only coated and uncoated PCL-TMA900 displayed no mineralisation as expected. The mice assessed displayed differing responses in bone formation capacity, which may be due to the mouse itself or likely due to differences in quantities of Laponite or BMP-2 attached to the scaffold. If the BMP-2 is dry and unbound on the surface of the scaffold this may have led to a burst release when surrounded by tissue giving fast, significant bone formation. Discrete variations in bone formation are not particularly useful in research, when looking for a clinically applicable material and the bone formation on the Laponite/BMP-2 coated PCL-TMA900 scaffolds illustrated the possibility of even greater bone formation as the scaffold is scaled-up.

The subsequent subcutaneous implantation study using Laponite/BMP-2 scaffolds stored for 24 hours i.e., ‘delayed Laponite/BMP-2’ scaffolds, showed unexpected results as the scaffolds were stored dry at RT (therefore the BMP-2 would not have been denatured). It is possible storage with a desiccant to prevent hydration of the coatings from air moisture or reducing storage temperature may maintain the bioactivity, or storage without touching a solid surface to reduce BMP-2 transfer from the scaffold. A single trial scaffold coated 6 months previously was implanted in mouse 4, to determine if long term storage of Laponite/BMP-2 coated materials could be an option, however, no bone formation was observed. In mice 1 and 2, the volume of bone formed on 24-hour dry delayed use coated scaffolds was approximately 33% and 12.5% of that formed on the immediately implanted coated scaffolds respectively. This delayed use of BMP-2 appears to have become active in mouse 3 after several weeks, as this bone formation was only visualised and quantifiable on *ex vivo* µCT scans at week 8. This leads to the speculation that the BMP-2 was initially on the surface of the scaffold, but the BMP-2 was also taken up by the rehydrated Laponite film during the 24-hour rehydration and macrophages may release the BMP-2 over time, however this would require further examination to test this theory. This mechanism of action of providing a local burst release followed by sustained release of BMP-2 would be ideal in bone tissue engineering.

The Laponite/BMP-2 coated scaffolds showed variation of bone volume formed in the subcutaneous implantation studies over the 8-week study periods. This may be due to aggregation of the Laponite particles due to gravity and therefore non-uniform coating of the hydrophobic PCL-TMA900 material in these mice. However, the mice which responded in the second study had greater mineralisation than in the first PCL-TMA900 Laponite/BMP-2 coated scaffold study and so variability does exist with this coating method, however, with time the active BMP-2 became available and sustained, gradual bone formation occurred *in vivo*.

In future work, optimisation of the Laponite and BMP-2 coatings to allow scaffold saturation with active BMP-2 will be important. The mass of BMP-2 held by the Laponite on solid polymer scaffolds and therefore delivered to the fracture site, will depend on the scaffold surface area, pore size, consistent uniform coating of the material, total mass of adherent Laponite and the saturation concentration/mass of BMP-2 within the Laponite. Optimising the BMP-2 held on the scaffold could be investigated using BMP-2 when the Laponite coating is a liquid or using the Laponite and BMP-2 mixed in solution to coat the scaffolds in one step. This proposed study could be robustly assessed in a murine subcutaneous implantation model as host macrophages will be important in releasing the BMP-2 from the Laponite to induce a cellular osteogenic response.

## 5. Conclusion

The current studies demonstrate the development of a biocompatible, nanoclay-coated, bioactive scaffold; Laponite and BMP-2 coating on PCL-TMA900 polymer. The absorption of BMP-2 to the Laponite proved consistent, with retained bioactivity and critically will be scalable. Due to the encouraging *in vitro* results, the Laponite/BMP-2 coating was found to be successful in producing heterotopic bone in the murine subcutaneous implantation model. The Laponite/BMP-2 coated scaffolds showed consistent bone formation, albeit, with variation in response time and volume of bone formed on the scaffold, with potential scope of use as a ready to use clinical product. This Laponite/BMP-2 coating has shown clear efficacy *in vitro* and *in vivo*, therefore, use of dry Laponite/BMP-2 coated PCL-TMA900 scaffolds in an osseous defect site would be the next stage in creating a clinically applicable scaffold for bone repair.

## Supporting information

Supplemental information Bioactive Laponite coated scaffolds

## Credit author statement

Øvrebø, Ø., Callens, S. J. P.: designed the scaffold shape using C.A.D software, Echalier, C. and Wojciechowski, J. P. synthesised the PCL-TMA and PCL-TMA900 resin, Wojciechowski, J. P. and Yang, T. prepared and printed the PCL-TMA and PCL-TMA900 scaffolds. Jayawarna, V. extrusion printed the PCL scaffolds and organised the EO sterilisation of scaffold materials. Marshall K. M. conducted the *in vitro* experiments, the CAM assay, murine subcutaneous implantation studies, data collection and analysis, histology and wrote the paper. Dawson J. I. supplied the Laponite. Kanczler J. M., Salmeron-Sanchez, M., Stevens, M. M., Dawson J. I. and Oreffo R. O. C. conceptualisation, funding acquisition, study supervision and editing of the manuscript.

## Data and materials availability

All data associated with this study are presented in the paper or the Supplementary Materials. All raw data is available on reasonable request from the corresponding authors.

## Acknowledgements

Research support for this study from the Biotechnology and Biological Sciences Research Council (BBSRC BB/P017711/1), the UK Regenerative Medicine Platform Acellular / Smart Materials – 3D Architecture (MR/R015651/1) and University of Southampton is gratefully acknowledged as well as many useful discussions with Dr Roxanna Ramnarine Sanchez and current members of the Bone and Joint Research Group in Southampton, UK. Dr Katie Dexter (University of Southampton, Biomedical Imaging Unit) is acknowledged for advice and expertise in µCT scanning of mice. For the purpose of open access, the author has applied a ‘Creative Commons Attribution (CC BY) license to any Author Accepted Manuscript version arising.

## Appendix A. Supplementary data

Supplementary data to this article can be found online.

