## Supplemental information Bioactive Laponite coated scaffolds for "Bioactive coatings on 3D printed scaffolds for bone regeneration: Use of Laponite™ to deliver BMP-2 for bone tissue engineering – progression through *in vitro*, chorioallantoic membrane assay and murine subcutaneous model validation"

### Supplementary information

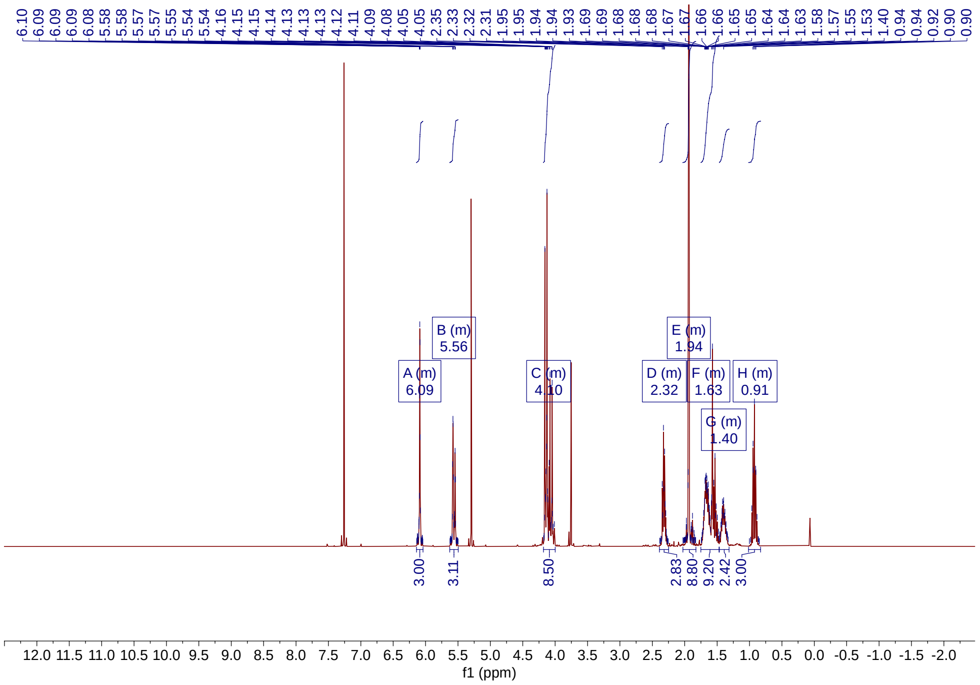


**S. Figure 1.** $^{1}H$ NMR spectra of poly(caprolactone)-trimethacrylate (PCL-TMA300) in $CDCl_3$. Comparison of the vinyl protons (d = 5.56, 6.09 ppm) integration against the propyl-$CH_3$ protons (d = 0.91 ppm) indicates a degree of functionalisation > 95%.

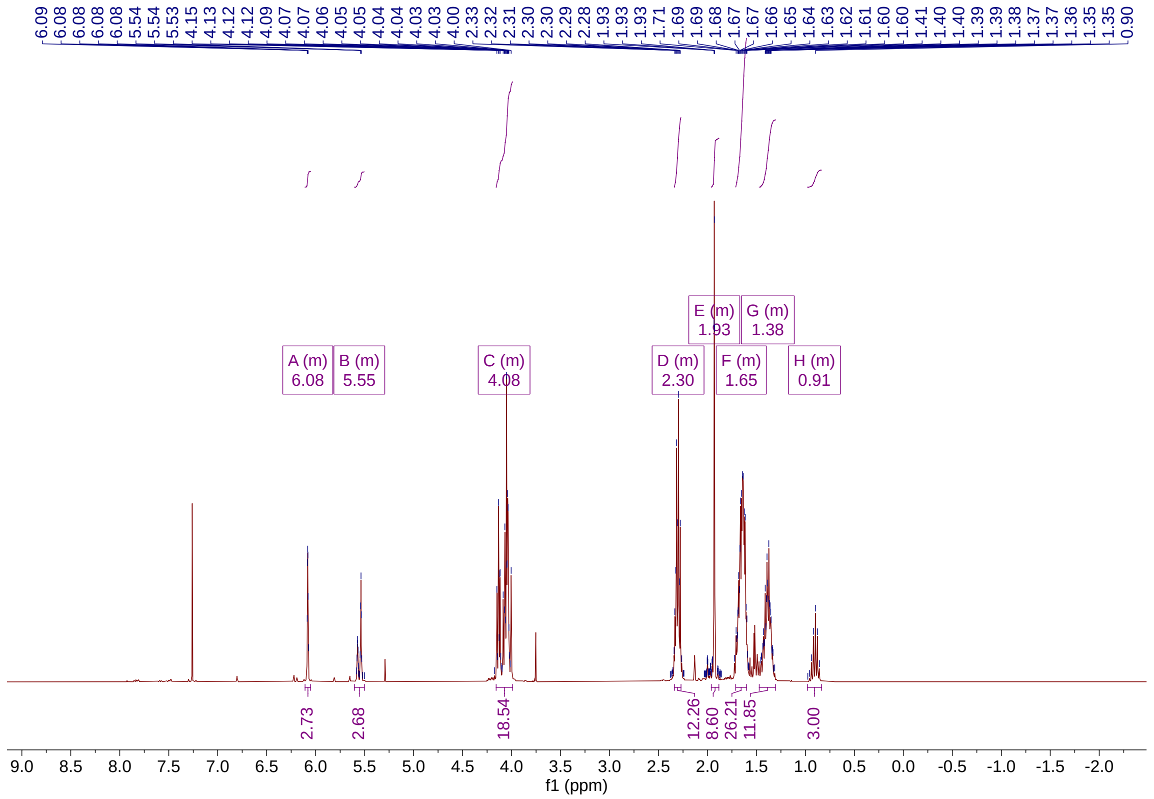


**S. Figure 2.** [1]H NMR spectra of poly(caprolactone)-trimethacrylate (PCL-TMA900) in $CDCl_3$. Comparison of the vinyl protons (d = 5.55, 6.08 ppm) integration against the propyl-$CH_3$ protons (d = 0.91 ppm) indicates a degree of functionalisation > 90%.

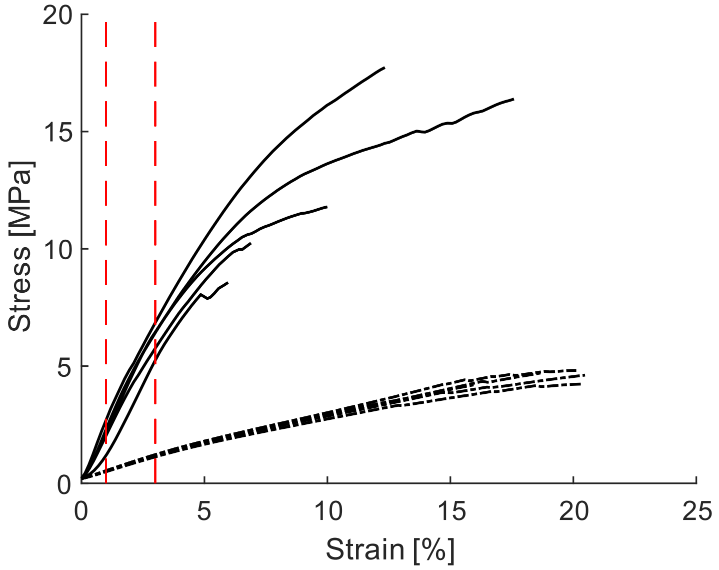


**S. Figure 3.** Compressive testing of the 3D printed PCL-TMA300 (–) and PCL-TMA900 (- -) unit cells (n = 5). The effective elastic modulus was determined between 1 – 3% strain.

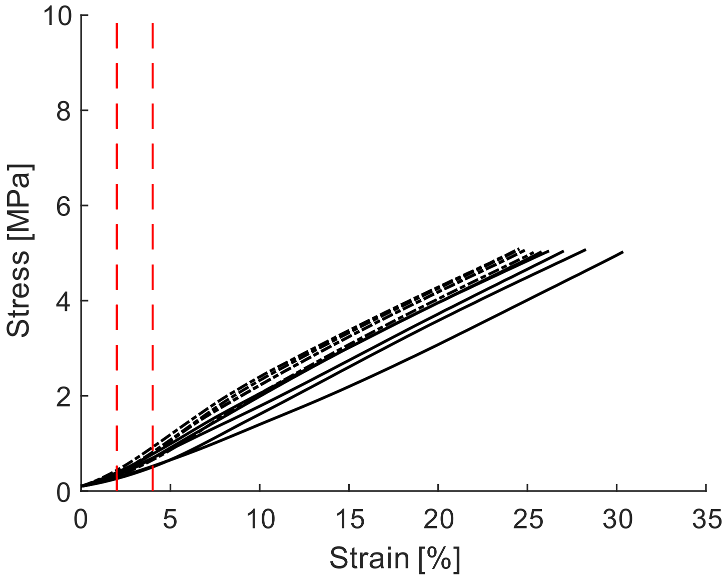


**S. Figure 4.** Compressive testing of 3D printed PCL-TMA900 following protocols from ISO 604. Samples were tested perpendicular (- -) and parallel (–) to the build axis (n = 5). The elastic modulus was determined between 2 – 4% strain.

**S. Table 1.** Summary of the compressive testing results on the 3D printed PCL-TMA300 and PCL-TMA900 unit cells.

| Sample | Effective E (MPa) | Average E (MPa) |
|---|---|---|
| **PCL-TMA 300** | | 203.7 ± 11 MPa |
| 1 | 205.7 | |
| 2 | 207.7 | |
| 3 | 186.2 | |
| 4 | 202.8 | |
| 5 | 216.3 | |
| **PCL-TMA 900** | | 33.9 ± 1.4 MPa |
| 1 | 34.2 | |
| 2 | 34.7 | |
| 3 | 35.2 | |
| 4 | 31.6 | |
| 5 | 33.7 | |

**S. Table 2.** Summary of the ISO 604 compressive testing results for 3D printed PCL-TMA900.

| Sample | E (MPa) | Average E (MPa) |
|---|---|---|
| **Perpendicular** | | 22.7 ± 2.6 MPa |
| 1 | 22.6 | |
| 2 | 22.2 | |
| 3 | 17.2 | |
| 4 | 24.2 | |
| 5 | 22.4 | |
| **Parallel** | | 15.6 ± 3.5 MPa |
| 1 | 18.2 | |
| 2 | 12.1 | |
| 3 | 16.6 | |
| 4 | 11.7 | |
| 5 | 19.3 | |

#### *Method for suspending Laponite and $dH_2O$ immersed PCL-TMA900 scaffolds to dry*

PCL-TMA900 and PCL scaffolds (for incubation time experiments in **Supplementary Figure 6**) were suspended using suture material through the pores of each scaffold to allow them to dry without touching a solid surface, to prevent transfer removal of the Laponite or $dH_2O$ from the scaffold surface (**Supplementary Figure 5**).

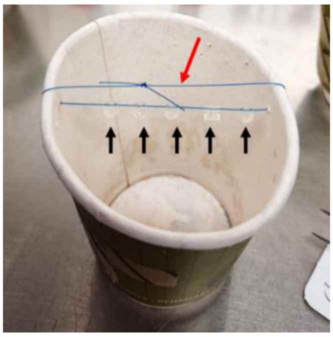

**S. Figure 5**: PCL-TMA900 and PCL scaffolds were suspended by suture material to dry. The PCL-TMA900 scaffolds shown (black arrows) were spaced apart, held on sterile nylon suture material (red arrow) tied through the sterilise plastic/paper cup to allow drying with the minimum scaffold area touching a surface.

#### *Assessment of the effect of incubation time in BMP-2 solution on osteogenic differentiation of C2C12 cells evidenced by ALP staining*

Circular scaffolds were designed using BioCAD software and fabricated with an extrusion-based 3D printer (RegenHU, Switzerland). Medical grade polycaprolactone (PCL) was loaded into a cartridge and heated to 70 °C for 1 hour until the polymer melted and stabilized. A 21 gauge stainless steel nozzle was used to extrude PCL strands under a pressure of 2 bars and a printing speed of 3 mm/second. Eight circular PCL scaffolds (5 mm diameter × 0.8 mm height) were added to 1% (w/w) Laponite in a 2 mL eppendorf, while nine scaffolds were placed in sterile $dH_2O$ in a 2 mL eppendorf. The scaffolds were left at RT for 1 hour and then suspended to dry for 2 hours, with the Laponite remaining and water evaporating leaving the scaffolds uncoated. The dry Laponite coated or uncoated scaffolds were submerged separately in BMP-2/PBS solution (5 µg/mL, 1 mL) in 1.5 mL low-protein binding eppendorf tubes to adhere BMP-2 to the surface of the scaffold and left at RT for the appropriate duration i.e., 5 days, 1 day, 1 hour. The scaffolds were removed and placed in a 24 well plate to dry for approximately 2 hours and placed in well plates for C2C12 cell seeding. C2C12 cells were suspended to $8x10^6$ per mL of DMEM/1% P/S/10% FCS (basal) media. A 25 µL aliquot of cell suspension ($2\times10^5$ C2C12 cells) was pipetted onto the PCL circular scaffolds and left to incubate at 37 °C for 30 minutes. The well was topped up with 975 µL of basal media or media containing 100 ng/mL or 5 µg/mL of BMP-2, for the positive control wells. Each condition was n=2 and the control conditions were n=1. The scaffolds were moved to a new 24 well plate and the appropriate fresh media added and not refreshed during culture. ALP staining was performed on day 5.

The ALP expression was more consistent at 1 and 5 days, with very faint staining after 1 hour in both $dH_2O$ treated and Laponite coated groups. The ALP expression after 5 days and 1 day was similar with obvious red staining, which was more intense in the Laponite coated group. The positive controls showed uniform ALP expression at 5 µg/mL of BMP-2 in the media and less ALP expression when 100 ng/mL was used. The negative controls displayed negligible ALP expression (**Supplementary Figure 6**).

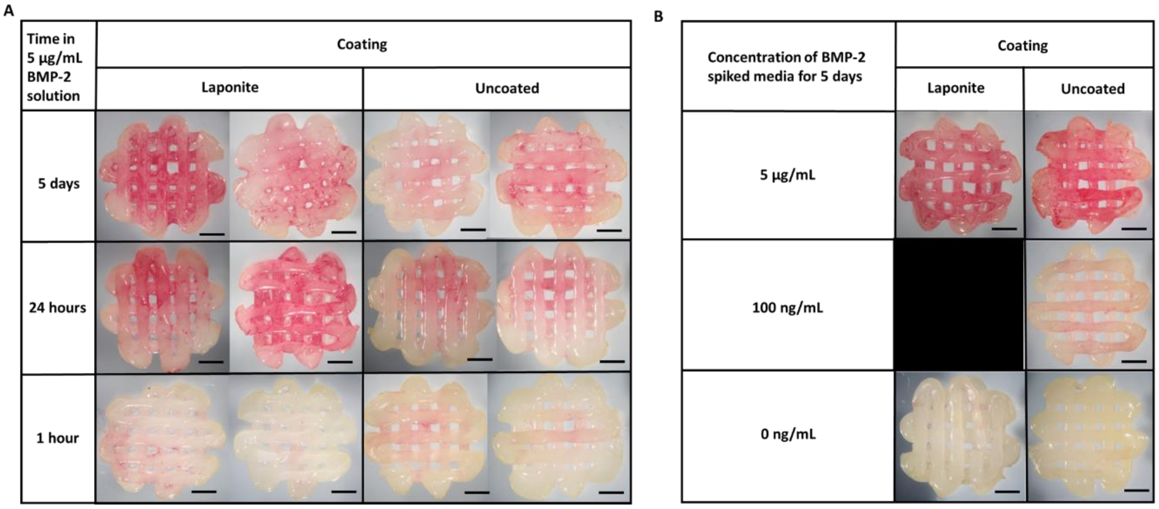


**S. Figure 6:** The effect of immersion time in BMP-2 solution. The ALP expression was most pronounced in the Laponite coated group and there was similar ALP expression at 5 days and 1 day timepoints, however there was less ALP expression at 1 hour. The positive controls showed similar ALP expression to the 1 day and 5 days Laponite coated groups, with intense red staining. The negative controls (no BMP-2 exposure) displayed negligible ALP expression evidenced by no staining observed. N=1 or 2 as shown, scale bar 1mm.

### C2C12 differentiation assay to ensure osteogenic activity

A BMP-2 dilution assay was set up to ensure the C2C12 cells could respond to BMP-2 by producing a red stain due to ALP production. $5\times10^4$ C2C12 cells were seeded into 12 wells of a 24 well plate in 500 µL DMEM/1%PS/10%FCS. After 24 hours the media was changed to 1 mL of DMEM/1%PS/2%FCS with no BMP-2 or 100 ng/mL, 50 ng/mL, or 25 ng/mL BMP-2 in a serial dilution. After 3 days the media was replaced and ALP staining performed on day 4 to ensure the C2C12 cells could respond to BMP-2 prior to use on scaffolds. The 100 ng/mL BMP-2 showed the most marked ALP expression with less expression seen at 50 ng/mL or 25 ng/mL and no obvious stain when no BMP-2 was added, as expected (**Supplementary Figure 7**).

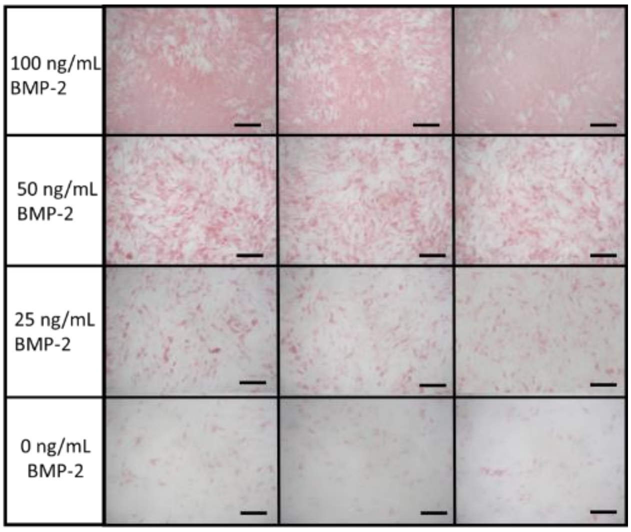


**S. Figure 7:** ALP staining of a BMP-2 dilution assay with C2C12 cells on TCP. This example aims to clarify that the cells were able to respond to BMP-2. A dose response to BMP-2 of reducing stain intensity with reducing BMP-2 concentration was observed, with negligible ALP expression at 0ng/ml BMP-2. N=3 for each condition, scale bar 500 µm.

### BMP-2 buffer solution formulation

The buffer solution for dilution of Medtronic InductOs® BMP-2 for use on collagen sponge discs was made by dissolving the following listed in **Supplementary Table 3** in $dH_2O$:

**S. Table 3:** InductOs® BMP-2 buffer solution components.

| **Reagent** | **Concentration** | **Quantity to make 500 mL** |
|---|---|---|
| Sucrose | 0.5% | 2500 mg |
| Glycine | 2.5% | 12500 mg |
| L-glutamic acid | 0.37% | 1850 mg |
| Sodium chloride | 0.01% | 50 mg |
| Polysorbate 80 | 0.01% | 50 mg |

### µCT scan settings used for *in vivo* and *ex vivo* imaging

The scan information for acquisition and reconstruction parameters for the *in vivo* and *ex vivo* µCT imaging is shown in **Supplementary Table 4**.

**S. Table 4:** Settings used for µCT scanning of mice and explanted samples in the subcutaneous implant studies.

| **Imaging reference** | **Samples *in vivo*** | ***Ex vivo* excised samples in sample holder** |
|---|---|---|
| Voltage (kVp) | 55 | 50 |
| Current (mA) | 0.17 | 0.21 |
| Exposure time (ms) | 75 | 65 |
| Filter (µm thickness) | Al 400+100 | Al 100 only |
| Scan angle (°) | 360 | 360 |
| Step angle (°) | 0.25 | 0.25 |
| Pixel size | 40 | 20 |
| Reconstruction voxel size ($\mu m^3$) | 40 | 20 |
| Frame averaging | 1 | 1 |
